# Super-resolution Molecular Map of Basal Foot Reveals Novel Cilium in Airway Multiciliated Cells

**DOI:** 10.1101/487330

**Authors:** Quynh P.H. Nguyen, Zhen Liu, Rashmi Nanjundappa, Alexandre Megherbi, Nathalie Delgehyr, Hong Ouyang, Lorna Zlock, Etienne Coyaud, Estelle Laurent, Sharon Dell, Walter Finkbeiner, Theo Moraes, Brian Raught, Kirk Czymmek, Alice Munier, Moe R. Mahjoub, Vito Mennella

## Abstract

Motile cilia are beating machines that play a critical role in airway defense. During airway cell differentiation, hundreds of motile cilia are templated from basal bodies that extend a basal foot, an appendage that links motile cilia together to ensure beating coordination. This assembly has thus far escaped structural analysis because its size is below the resolution limit. Here, we determine the molecular architecture and identify basal foot proteins using a super-resolution-driven approach. Quantitative super-resolution image analysis shows that the basal foot is organized in three main regions linked by elongated coiled-coil proteins. FIB-SEM tomography and comparative super-resolution mapping of basal feet reveal that, among hundreds of motile cilia of an airway cell, a hybrid cilium with features of primary and motile cilia is harbored. The hybrid cilium is conserved in mammalian multiciliated cells and originates from parental centrioles. We further demonstrate that this novel cilium is a signalling centre whose cellular position is dependent on flow.

## Introduction

Motile cilia are beating machines that generate the propulsive force required for mucociliary clearance, thereby protecting the airways from recurrent infections and environmental pollutants^1,2^. To beat in coordination, motile cilia rely on the basal foot, a triangular structure attached to the basal body on one end and to the microtubule cytoskeleton on the other, thereby linking hundreds of motile cilia together in a network^3,4,5^. The hundreds of motile cilia on the surface of an airway multiciliated cell are thought to be similar to each other and templated from identical basal bodies each presenting at their base one basal foot^6,7^ pointing toward the direction of ciliary beating^8,9^ —a phenomenon termed rotational polarity^10,11^.

In airway cells, loss of the basal foot results in disruption of the microtubule apical network, irreversible disorientation of basal bodies and lack of motile cilia coordination^4^. In mice, loss of the basal foot leads to respiratory manifestations indicative of Primary Ciliary Dyskinesia (PCD)^4,12^, an autosomal recessive disease characterized by chronic airway infections, and frequently associated with hearing loss, male infertility, hydrocephalus and heterotaxy, which can lead to lung collapse and death in mid adulthood^13,14,1,2,15,16^. Despite the basal foot’s critical role in airway physiology and multiciliated cell function, its molecular organization remains to be elucidated.

In cells protruding a primary cilium, the basal foot—along with centrosomal proximal end proteins—keeps the primary cilium submerged by linking the basal body to the Golgi network thereby avoiding ectopic Shh-signalling^17,18^. Differently from motile cilia, the basal foot of the primary cilium is present in multiple copies per basal body and is thought to originate from nine (or less depending on the cell type) subdistal appendages, mother centriole-associated structures contributing to interphase microtubule organization^21,20,8^ (Fig. 1a). In mammalian cells, subdistal appendages appear in electron microscopy (EM) micrographs as thin, conical-shaped structures with a round tip connected to the centrosomal barrel by two microtubule triplets^22,21,23,24^. Less is known about the basal foot structure and composition in primary cilia, and no consensus has been reached on its nomenclature: this assembly has been named differently depending on the study and cell type (e.g. satellite arms^21^, basal feet^25^ or subdistal appendages^17^).

Several proteins have been assigned to the basal foot and subdistal appendages in mammalian cells through conventional fluorescence microscopy and immuno-EM (Ninein (NIN)^26^, ODF2/Cenexin^4,27,28^, CC2D2A^29^, CEP170^30^, Galactin-3^3^, ε-Tubulin^31^, Centriolin (CNTRL)^32^, Trichoplein (TCHP)^33^, CEP128^17^, CEP19^34^, CCDC120 and CCDC68^35^). ODF2/Cenexin is a fibrillar protein related to the intermediate filament (IF) superfamily that plays a critical role in basal foot assembly since lack of basal foot-specific ODF2/Cenexin isoform results in loss of the entire structure^4,5,12,36^. ODF2/Cenexin interacts with TCHP, an IF-binding protein implicated in the recruitment of NIN to subdistal appendages^33^. NIN and CEP170 have been implicated in microtubule anchoring and nucleation functions^38,39,40^. Recently, CEP19 and CC2D2A have been assigned to subdistal appendages with the latter shown to play a critical role in their assembly^34,29^.

Despite the information has accumulated on individual proteins, to date a comprehensive and quantitative view of the molecular architecture of the basal foot is still lacking. Moreover, it remains unknown how the basal foot’s organization changes in different cilia or in subdistal appendages to accommodate its specific functions.

Here, we resolve the structure of the basal foot in cilia *in situ* using super-resolution microscopy revealing an architecture composed of three regions linked by elongated proteins, which is partly conserved in different types of cilia. Unexpectedly, our super-resolution analysis reveals a novel “hybrid” cilium in multiciliated cells characterized by a basal body with multiple basal feet. The hybrid cilium originates from parental centrioles and presents structural features of a motile cilium. Functional analysis using airway cells from healthy individuals and patients with immotile cilia syndrome suggests that the hybrid cilium position is dependent on flow generated by the surrounding motile cilia. Altogether, our data show that not all motile cilia are identical beating machines in a multiciliated cell. Furthermore, they provide evidence of a novel sensing mechanism in multiciliated epithelia.

## Results

### Super-resolution Microscopy and BioID Reveals Structural Organization and Novel Components of the Basal Foot

To determine the molecular architecture of the basal foot *in situ*, we first focused on primary cilia from immortalized Retinal Pigment Epithelia 1 (hTERT-RPE1) cells, a cellular model characterized by robust ciliation and homogenous ciliary structure. We reasoned that 3DSIM microscopy resolution power (∼125 nm in x/y and ∼250 nm in z axis)^41,42,43^ was sufficient to assign proteins to the basal foot and/or to other ciliary regions (Fig. 1a). To test this, we examined the distribution of NIN, a basal foot protein reported to have different sub-populations at the basal body^44^. Using 3DSIM, we clearly distinguished three sub-populations of NIN: one at the proximal end of the basal body, one at the daughter centriole, and a third at the basal foot, which extends laterally at the distal end of the basal body (Fig. 1b). We then quantitatively mapped the position of all reported basal foot/subdistal appendage proteins relative to the centre of the basal body measured by polyglutamylated-Tubulin, a modification of centriolar microtubules that is a proxy for the outer diameter of centrioles (∼200 nm; Fig. 1c)^45^. Since 3DSIM resolution is maximum in the x/y plane, in-plane end-on and side-views were selected for measurement from hundreds of micrographs with basal foot proteins labeled with 488-conjugated secondary antibodies to ensure highest resolution power (Fig. 1c, Sup. Fig. S1).

**Figure 1.**
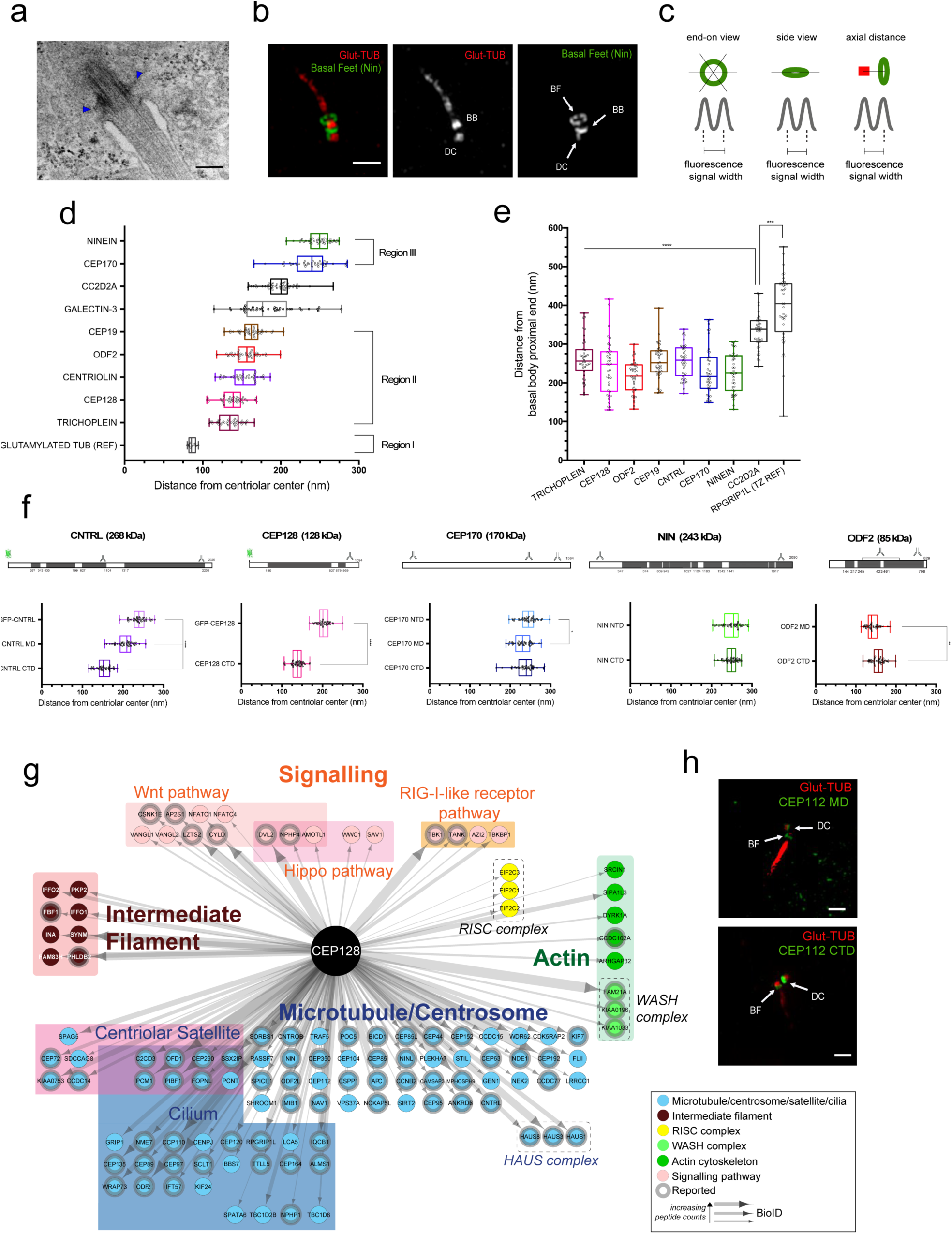
3D-SIM and BioID reveal the molecular architecture and a novel component of basal foot in primary cilia. **(a)** Representative TEM micrograph of a primary cilium and its basal feet (blue arrowheads) in immortalized RPE-1 cells. Scale bar represents 200 nm. **(b)** 2D projection micrograph of 3DSIM volume of a primary cilia in RPE-1 cell stained with anti-Ninein (NIN, green) and anti-glutamylated tubulin (Glut-TUB, red) antibodies, showing the three known subpopulations of the protein: at the proximal ends of basal body, daughter centriole and at the basal feet (see arrows). Scale bar represents 1 μm. **(c)** Cartoon depiction of the strategy used to measure radial and axial distance of basal foot proteins (green) in primary cilia. The radial distance, or distance from the centriolar center, was calculated either from end-on view by dividing each of the ring diameter measurements by two (left) or from side views by measuring the lateral distance of basal foot proteins positioned across the basal body (middle). Axial distance was measured relative to the basal body proximal end (red). **(d)** Box plot of radial distances of basal foot proteins in primary cilia of RPE-1 cells (n=40). Region assignment of proteins was based on statistical analysis by one-way ANOVA using Tukey’s multiple comparison test. Distance measurements of proteins not significantly different from each other were grouped into the same region. **(e)** Box plot of axial distances of basal foot proteins in primary cilia from RPE-1 cells (n=40). RPGRIP1L was used to label the transition zone. Statistical analysis was done by one-way ANOVA with Tukey’s multiple comparison test. **(f)** Top: linear maps representing protein polypeptide sequences showing the regions recognized by antibodies and the position of GFP insertion. Bottom: Box plot of radial distributions of CNTRL, CEP128, CEP170, NIN and ODF2 in primary cilia of RPE-1 cells using domain specific antibodies. Statistical analyses were conducted using Welch’s t-test (for pair-wise comparisons) and Tukey’s test (for multiple comparisons). Unless indicated otherwise, the differences are not significant. **(g)** Diagram showing proteins identified via BioID in close proximity to CEP128-BirA* in ciliated HEK-293 cells. Arrow thickness is proportional to the number of peptides detected. **(h)** 2D projection micrographs of 3D-SIM volume of a primary cilium in RPE-1 cell stained with anti-CEP112 (green) antibodies labelling middle domain (MD, left) and C-terminal domain (CTD, right), and anti-Glut-TUB (red) antibody, showing two distinct subpopulations of the protein: at the proximal ends of daughter centriole and at the basal feet (see arrows). Scale bars represent 1 μm.

Notably, 3DSIM mapping shows that basal foot proteins are clustered into spatially separated regions (Fig. 1d, Table S1). NIN and CEP170 are the most distant from the centriole center (248±16 nm and 237±25 nm, respectively), consistently with their association with microtubules^26,38,39^. Therefore, this region was termed the microtubule-anchoring region or region III. Most basal foot proteins are clustered with ODF2, a component critical for basal foot assembly^4,5^ (ODF2: 155±16 nm; CEP128: 139±15 nm; CEP19: 166±15 nm; and CNTRL: 153±18 nm; TCHP: 135±15 nm; Galactin-3 shows a broad distribution centred around this region (185±36 nm). Since this intermediate region contains ODF2, it was termed the scaffolding region or region II. Interestingly, our imaging map shows a gap where the basal body connects to the basal feet, a region that was termed the basal body anchoring region or region I. Among the proteins previously assigned to the basal foot/subdistal appendage, ε-tubulin, CCDC120, CCDC68 could not be reliably detected at the basal foot with available commercial antibodies, while TCHP, a protein thought to be associated only with subdistal appendages^31,35^, was located to the basal foot in primary cilia. To correctly assign proteins to the basal foot, we then measured the position of basal foot proteins along the axoneme relative to the proximal end of the basal body (Fig. 1c, e). As expected, most basal foot proteins were distributed in the same axial region (209-284 nm), with the exception of CC2D2A, whose c-termini was located significantly above the basal foot (333±22 nm) and below the transition zone labeled with RPGRIP1L, a *bona fide* transition zone protein that is part of the Y-links (393± 90 nm; Fig. 1e)^46^. This suggests that CC2D2A is located not exclusively within the basal foot region.

To ensure basal foot structural integrity, we hypothesized that some proteins must be connecting different regions of this supramolecular assembly together as molecular linkers either in the form of pearls on a string and/or elongated proteins^47,48,49^. Since most basal foot proteins showed similar distribution variances, linkers were likely to be high-molecular weight, elongated coiled-coil proteins, similar to centrosomal proteins of the pericentriolar material^47^. We then used antibodies and GFP-fusion proteins labeling different protein domains to identify their position within the basal foot (Fig. 1f). Notably, CNTRL was found to link regions II and III by extending over a distance of ∼75 nm (CNTRL C-Terminal Domain (CTD): 153±18 nm; GFP-CNTRL: 240±19 nm; Fig. 1g). CEP128 also showed an extended organization, but not as far from the basal body as CNTRL (CEP128 CTD: 139±15 nm; GFP-CEP128: 203±16 nm; Fig. 1g). Interestingly, NIN also showed an elongated distribution looping back toward region II (Sup. Fig. S2, Sup. Table S2). In contrast, CEP170 did not appear extended consistently with its lack of coiled-coil domains (CEP170 CTD: 237±25 nm; Middle Domain (MD): 231±22 nm; N-terminal Domain (NTD): 244±22 nm, Fig. 1g). Altogether, our data rule out a model where a single basal foot component spans the whole structure acting as a scaffold for the recruitment of other components. It shows instead that the basal foot is organized in distinct structural regions: region III is the microtubule-anchoring/nucleation region made of proteins CEP170 and NIN corresponding to the basal foot cap; region II consists of the majority of known basal foot proteins (TCHP, CEP128, ODF2, CEP19 and CNTRL) and region I anchors the basal foot to the basal body, though its composition is not yet well characterized. Subdistal appendages show a similar architecture to the basal foot of primary cilia, suggesting that their structure remains largely conserved during the transition from centrioles to basal bodies in primary cilia despite the change in the number of appendages and/or possible changes in composition (Sup. Fig. S3, Sup. Table S3).

Since our map showed few basal foot proteins in the region closer to the basal body, this suggested that new basal foot proteins might have yet to be identified and if so, they should be located in close proximity to CEP128, the coiled-coil protein nearest to the basal body. To test this possibility, we mined BioID data from datasets that used BirA* fused to the N-terminus of CEP128 (BirA*-CEP128), the furthest CEP128 domain from the basal body (Fig. 1f)^34^, and performed Bio-ID experiments using CEP128-BirA*, where BirA* is fused to C-terminus of CEP128, the closest domain of CEP128 to the basal body (Fig. 1g). CEP128 proximity map included many centrosomal, ciliary and satellite proteins as expected, but also components of different cytoskeletal structures including actin and intermediate filaments. To identify novel components of the basal foot, we then followed up on CEP112, a protein previously assigned to the centrosome^50^ that is enriched in coiled-coil domains and was identified in BioID experiments in close proximity to both centriolar marker CEP135 and to basal foot region II marker CNTRL. Immunolabeling of CEP112 with two antibodies raised against different epitopes of the protein showed a localization pattern consistent with that of a basal foot protein located close to the basal body centre (CTD: 128±13 nm; MD: 142±14 nm), with a second population at the daughter centriole (Fig. 1h, Sup. Fig. S4).

### Super-resolution Microscopy of the Basal Foot in Motile Cilia

We next asked whether the architecture of the basal foot was conserved in motile cilia. Since the basal foot plays an important structural role linking motile cilia together to ensure proper beating coordination, it is likely organized differently than in primary cilia^11,4,5^. TEM micrographs of basal foot from sections of human airway multiciliated cells suggest both structural similarities and differences (Fig. 2a). The basal foot appears by EM as a conical structure (h= ∼130 nm and w= ∼200 nm) attached to the basal body by three microtubule triplets with several electron-dense regions, including a bulky domain at the tip, the basal foot cap^3,21^. Similarly to primary cilia, the basal foot cap has a round structure with anchored microtubules (Fig. 2a, asterisk) and it appears connected with the central region of the basal foot by fibrils (Fig. 2a, blue arrrowhead). Differently from primary cilia, the basal foot is wrapped by two spherical structures symmetrically positioned close to and on the side of the tip and it is connected to the basal body by arches originating from three axonemal microtubules triplets (Fig. 2a, red arrowheads).

**Figure 2.**
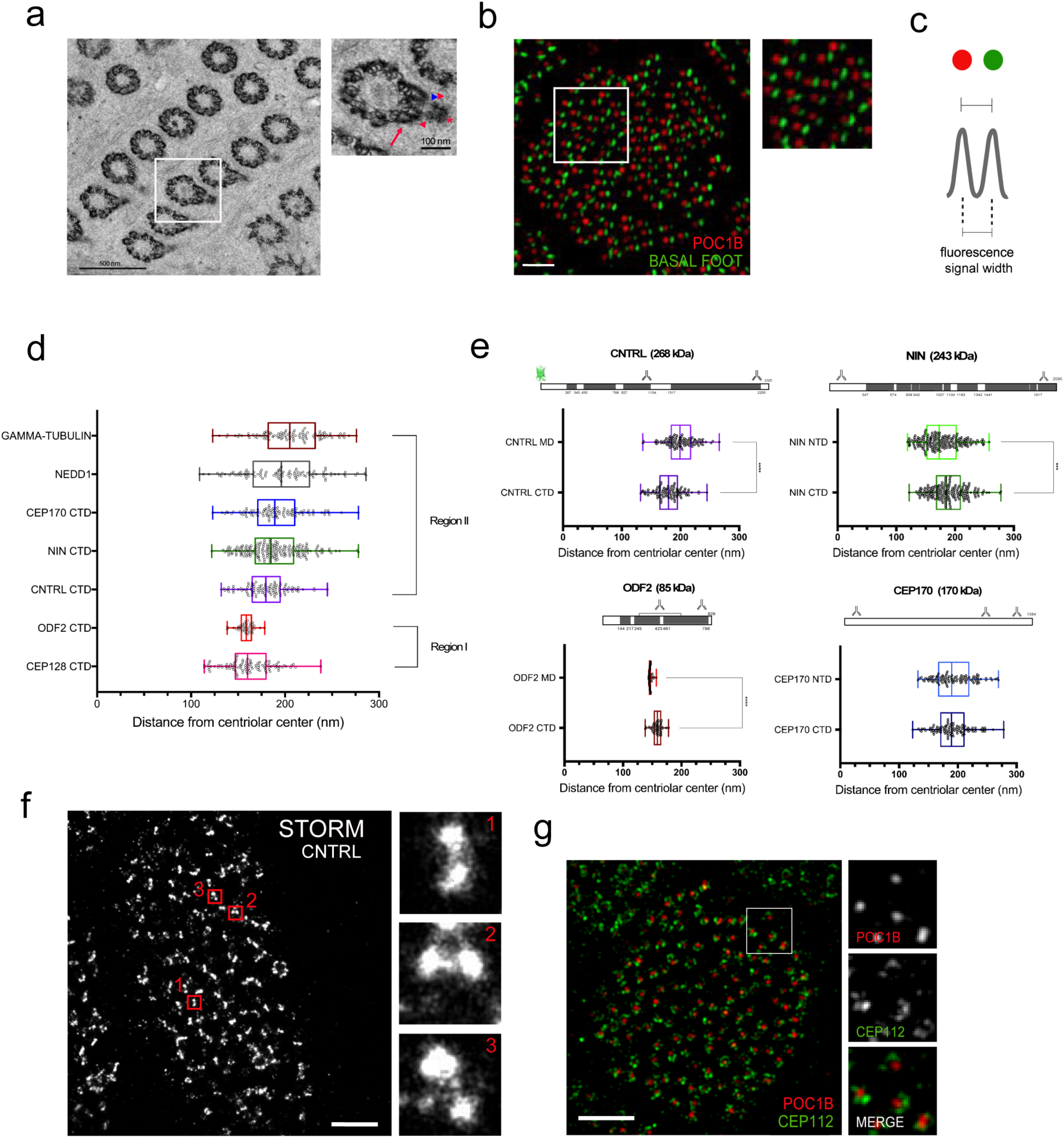
The conserved and distinct architectural features of basal foot in motile cilia. **(a)** Left: Representative TEM micrograph from a cross section of a human airway multiciliated cell. Scale bar represents 200 nm. Right: High-magnification view of the boxed area. Note three main electron-dense regions of the basal foot. Red asterisk denotes basal cap region, red arrowheads the spherical symmetrical structures, blue arrowhead the fibril region and red arrow the arch-region. Scale bar represents 100 nm. **(b)** Left: 2D projection micrograph of 3D-SIM volume of an airway epithelial multiciliated cell (end-on view) labeled with an antibody recognizing a basal body protein (POC1B, red) and a basal foot protein (CNTRL, green). Scale bar represents 1 μm. Right: High-magnification view of boxed area. **(c)** Cartoon depiction of the strategy used to measure radial distance of basal foot proteins (green) in motile cilia of human airway multiciliated cells using end-on view. Radial distance was measured relative to the basal body center (red). **(d)** Box plot of radial distributions of basal foot proteins in human airway multiciliated cells (n=80). Region assignment was done based on one-way ANOVA with Tukey’s multiple comparison test. Proteins whose distances were not significantly different were grouped into the same region. **(e)** Top: linear maps representing protein polypeptide sequences showing the regions recognized by antibodies. Bottom: Box plot of radial distributions of CNTRL, NIN, ODF2 and CEP170 in motile cilia of human airway multiciliated cells using domain specific antibodies. (n=80). Statistical analyses were conducted using Welch’s t-test for pair-wise comparisons. **(f)** Left: 2D-STORM micrograph of human airway multiciliated cells labeled with anti-CNTRL antibody. Scale bar represents 1 μm. Right: High-magnification views of boxed areas. **(g)** 2D projection micrograph of 3D-SIM volume of an airway epithelial multiciliated cell (end-on view) labeled with an antibody recognizing a basal body protein (POC1B, red) and CEP112 CTD antibody. Scale bar represents 2 μm.

To assign basal foot proteins *in situ* in motile cilia, we used POC1B a component of the basal body as a reference marker (Fig. 2b)^51,52^. Since in motile cilia there is only one basal foot per basal body, the pattern of basal foot proteins by 3DSIM appeared as a diffraction-limited spot, and not as a ring-pattern as observed in primary cilia (Fig. 1b, c). Taken together, the molecular map data show that basal foot molecular architecture is only partly conserved between motile and primary cilia (Fig. 2d, e, Sup. Table S4). NIN and CEP170 are located at the basal foot tip together with γ-Tubulin and NEDD1, proteins which are part of the γTuRC microtubule nucleating complex ^53^. The position of γ-Tubulin and NEDD1 was measurable only in motile cilia since basal bodies are largely devoid of Pericentriolar Material, the protein network surrounding the centriolar core^54^. Region II components CEP128 (162±25 nm) and ODF2 (158±16 nm) in motile cilia showed similar distances from the centriole wall as in primary cilia, while CNTRL is located further away from the basal body centre (180±23 nm and 153±18 nm, respectively) suggesting a distinct organization of this bridging protein in motile cilia. Notably, STORM microscopy shows that CNTRL is distributed in two main populations at the basal foot, located on opposite side of its longitudinal axis (Fig. 2f). Comparison of the measurements from EM micrographs with the ones from super-resolution images suggests that CNTRL fluorescence is located where two electrondense spherical structures are detected in EM sections (distance of the basal body center to spherical structures by EM: 190±15 nm; CNTRL distance from basal body center by 3DSIM: 180±23 nm). In fully differentiated multiciliated cells, CEP112 was observed as a small arch on one side of the basal body in close proximity to Cep128 (d=165±28 nm, Fig.2g) or a more closed ring in cells that appear not completely mature suggesting a dynamic distribution of the protein during differentiation (Sup. Fig. 5a). Cep19, on the other hand, forms a complete ring more similar to distal appendages proteins throughout docking and motile cilia extension. Lastly, TCHP showed a complex filamentous distribution that did not allow accurate measurements (Sup. Fig. S5). Altogether, our data show that basal foot architecture in primary and motile cilia is composed of modular regions that are largely conserved away from the basal body, while different in the basal body attachment region.

### Super-resolution Reveals a Novel Type of Cilium in Airways Multiciliated Cells

The comparison of the super-resolution maps of basal foot in primary and motile cilia indicated that functionally different cilia have structural and numerical differences: there are multiple basal foot per basal body in primary cilia, and only one basal foot per basal body in motile cilia (Fig. 1b and Fig. 2b). In human multiciliated cells, it was therefore surprising to observe a ring pattern of basal foot proteins, since at the base of motile cilia only a single basal foot per basal body was thought to be present (Fig. 3a and Sup. Fig. S6). This arrangement suggested the presence of a basal body similar to the primary cilium one.

**Figure 3.**
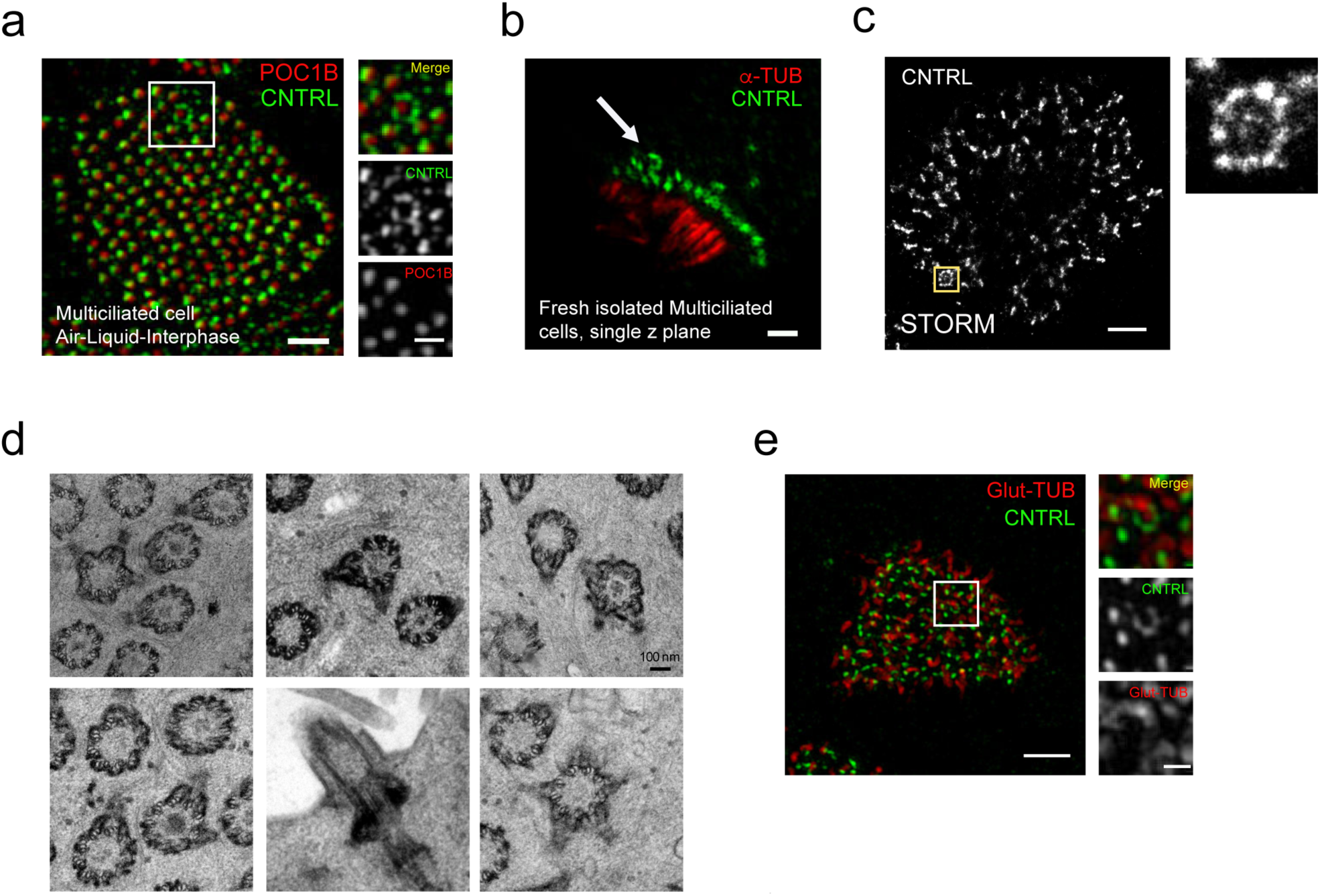
Super-resolution mapping of basal foot reveals a novel type of cilia in airway multiciliated cells. **(a)** Left: 2D projection micrograph of 3D-SIM volume of an airway multiciliated cell grown on Air-Liquid-Interface (ALI), labeled with anti-CNTRL (green) and anti-POC1B (red) antibodies. Note the ring-like pattern of CNTRL localization encircling the basal body labeled by POC1B. Right: High-magnification view of boxed area. Scale bars represent 1 μm (left) and 500 nm (right). **(b)** 2D projection micrograph of 3D-SIM volume of human airway multiciliated cells freshly isolated from healthy individual, labeled with anti-CNTRL (green) and anti-alpha-tubulin (red) antibodies, showing the presence of the basal body with multiple basal feet. Scale bar represents 1 μm. **(c)** Left: 2D-STORM micrograph of airway multiciliated cell labeled with anti-CNTRL antibody, showing a distinct ring-like distribution of CNTRL. Right: High-magnification view of boxed area. Scale bars represent 1 μm. **(d)** Collage of representative TEM micrographs showing basal bodies harboring multiple basal feet in human airway multiciliated cell. Scale bars represent 100 nm. **(e)** Left: 2D projection micrographs of 3DSIM volume of an airway multiciliated cell labeled with anti-CNTRL (green) and anti-Glut-TUB (red) antibodies. Note the axoneme emanating from ring-like structure labeled with CNTRL. Right: High-magnification view of boxed area with individual channels. Scale bars represent 2 μm (left) and 500 nm (right).

3DSIM and STORM micrographs from human multiciliated cells co-labeled with antibodies recognizing basal foot (CNTRL, CEP128) and basal body (POC1B) proteins demonstrated that the ring pattern resulted from a basal body with multiple basal feet and not from multiple basal bodies clustering together as in the compound cilia, a membrane-delimited structure made of multiple motile cilia clustered together and frequently found in airway cells after injury^55^ (Fig. 3a, c; Sup. Fig. S6). To further confirm its *in vivo* relevance and organization, we established its presence in freshly isolated human upper airway cells (Fig. 3b) and in TEM micrographs (Fig. 3d). Furthermore, we then used 3DSIM to verify that an axoneme is emerging from it therefore demonstrating that this special basal body templates a cilium (Fig. 3e).

### The Novel Cilium has Hybrid Features of Primary and Motile Cilia

Next, we examined the ultrastructure of this unique cilium using Focus Ion Beam-Scanning Electron Microscopy tomography (FIB-SEM). Analysis of tomogram sections (Fig. 4a-c, Sup. Movie 1, 2) confirmed that the basal body templates a bona fide cilium and demonstrated that its axoneme contains a central pair similar to the surrounding motile cilia (Fig. 4a, b and Sup. Video 1-3). 3DSIM micrographs of airway multiciliated cells labeled with anti-radial spoke head (RSPH4A) and nexin-dynein regulatory complex (GAS8) antibodies further demonstrated that this special cilium harbors proteins specific for the ciliary beating machinery (Fig. 4d)^56,57^. Altogether, our data show that in human multiciliated cells of the airway, not all the cilia are identical; rather, a single cilium per cell on average presents hybrid features of primary and motile cilia.

**Figure 4.**
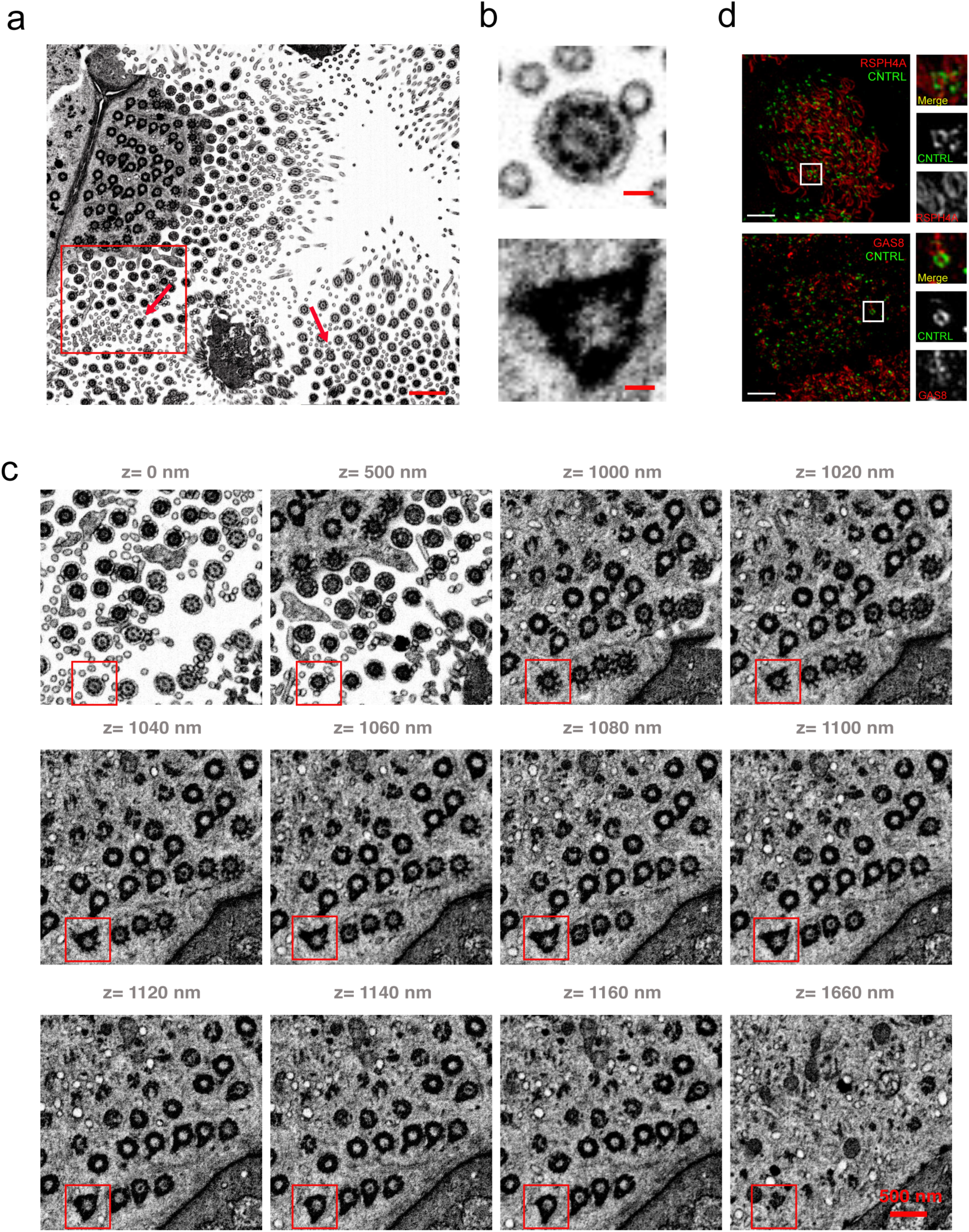
The novel cilium has hybrid features between primary and motile cilia. **(a)** Representative section from FIB-SEM tomogram of human multiciliated cells. Arrows indicate basal bodies with multiple basal feet. Scale bar represents 1 μm. **(b)** High-magnification view of boxed area in (a) at different z position of the tomogram from z=0 nm to z=1660 nm. Note the hybrid cilium axoneme and central pair (z=0 nm), transition fibers (z=1000-1040 nm), multiple basal feet (z=1160-1160 nm) and the absence of the endocytic pocket (z=1660 nm). Scale bar represents 500 nm. **(c)** A high-magnification view of boxed area in (b) highlighting the basal body with a central pair and multiple basal feet. Scale bar represents 100 nm. **(d)** 2D projection micrographs of 3D-SIM volume of human airway multiciliated cells (left), and high-magnification views of boxed areas (right), labeled with anti-CNTRL (green), anti-RSPH4A (red, top) and anti-GAS8 (red, bottom) antibodies. Scale bars represent 2 μm.

### Hybrid Cilium is Conserved in Different Mammalian Multiciliated Cells and Originates from Parental Centrioles

To determine if the hybrid cilium is conserved in other multiciliated cells and mammalian species, we examined multiciliated cells differentiated from progenitor basal cells isolated from adult mouse tracheal and ependymal tissue (Fig. 5a, b). Notably, when using 3DSIM microscopy and TEM we observed a basal body surrounded by multiple basal feet in these multiciliated cells demonstrating that the hybrid cilium is present in multiple tissues in mice and humans.

**Figure 5.**
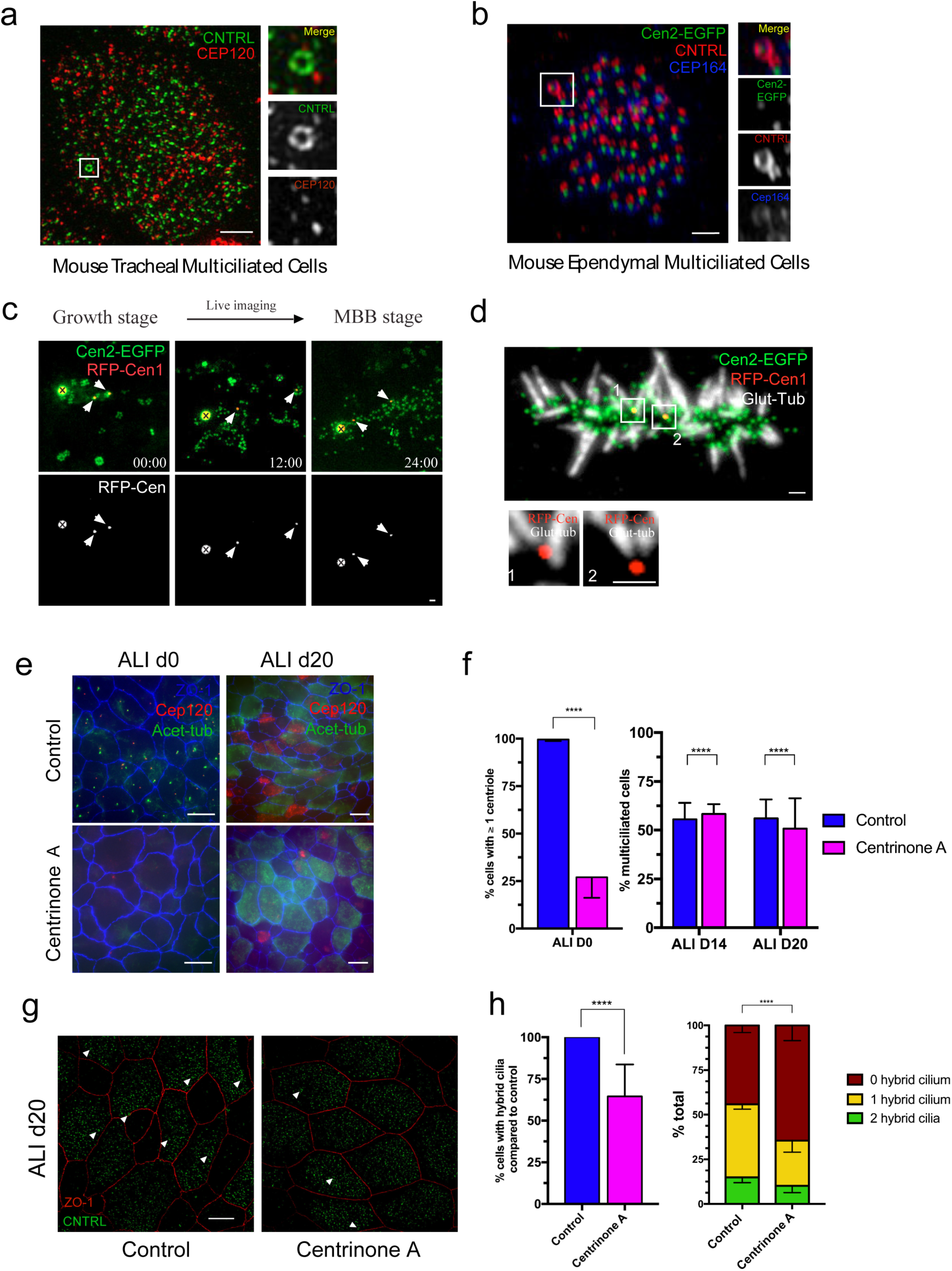
Hybrid cilium is a conserved feature of multiciliated cells and originates from parental centrioles. **(a)** Left: 2D projection micrograph of 3D-SIM volume of mouse tracheal multiciliated cell (ALI D20), labeled with anti-CNTRL (green) and anti-CEP120 (red) antibodies. Right: High-magnification view of boxed area with individual channels. Scale bar represents 2 μm. **(b)** Left: 2D projection micrograph of 3D-SIM volume of adult mouse ependymal multiciliated cells (P16), labeled with GFP-Centrin2, anti-CNTRL (red) and anti-CEP164 (blue) antibodies. Right: High-magnification view of boxed area labeled in left. Scale bar represents 1 μm. **(c)** Live imaging of TagRFP-Cen1 centrosomal centrioles during centriole amplification in primary cultured ependymal progenitors from Cen2-EFGP mice. Newly formed EGFP+ procentrioles are growing from deuterosomes and RFP+ centrosomal centrioles (00:00) before disengaging from their growing platforms (12:00) and gathering all together in the basal body patch (24:00). Arrowheads point to RFP+ centrosomal centrioles. A «x» sign marks centrin aggregates. **(d)** 2D projection micrograph from immunostaining experiment of primary cultured ependymal multiciliated cells labeled with antibody labeling glutamylated-tubulin (GT335) and RFP+ centrosomal centrioles that are retained in the basal body patch. Note cilia growing from centrosomal centrioles **(e)** 2D projection fluorescence micrograph of volumes of mouse tracheal multiciliated cells at ALI D0 (left) and ALI D20 (right), treated with DMSO control (top) or Centrinone A (bottom) and labeled with anti-acetylated tubulin (green), anti-CEP120 (red) and anti-ZO-1 (blue) antibodies. Scale bars represent 10 μm. **(f)** Bar graph showing percentage of cells with more than one centrioles at ALI D0 (left) and percentage of multiciliated cells at ALI D14 and D20 (right) in DMSO control (blue) or Centrinone A (pink); n>6000. Statistical analysis was done using Cochran-Mantel-Haenszel test. **(g)** 2D projection micrograph of 3D-SIM volume of mouse tracheal multiciliated cells at ALI D20 treated with DMSO control (left) or Centrinone A (right), labeled with anti-CNTRL (green) antibody. Arrowheads indicate CNTRL rings. Scale bar represents 5 μm. **(h)** Left: Bar graph representing percentage of cells with hybrid cilium in DMSO control (blue) and Centrinone A-treated (pink) cells, normalized to control condition; n>800. Statistical analysis was done using Cochran-Mantel-Haenszel test. Right: Bar graph representing percentage of cells with none (red), one (yellow) or two (green) hybrid cilia in ALI D20 mouse tracheal multiciliated cells treated with DMSO control (left) or Centrinone A (right); n>800.

The presence of multiple basal feet typical of primary cilia suggested that the hybrid cilium might be derived from the parental (mother) centriole, which first templates the primary cilium present during the early stages of differentiation of airway multiciliated cells, and then is resorbed before centriole amplification through the canonical and deuterosome pathway^58,59,7^. To first test whether the mother and daughter centrosomal centrioles are retained in differentiated multiciliated cells^58,60^, we performed a pulse-chase experiment which allowed us to label differentially centrosomal centrioles from newly formed basal bodies in mouse cultured ependymal cells (Supplemental Fig. S7). Time-lapse monitoring of RFP-Cen1 centrosomal centrioles in cells from Cen2-EGFP transgenic mice^61^ showed that centrosomal centrioles are retained within the newly formed basal body patch (Fig. 5c) and are capable of growing cilia (Fig. 5d). To further test that they indeed provide a template for the hybrid cilium, we used Centrinone, a small molecule drug blocking canonical centriole duplication by inhibiting Plk4, a kinase critical for the early stages of centriole duplication^62^. As expected, basal cells isolated from mouse trachea treated with Centrinone showed a reduction in the number of centrioles before airway cells differentiation into an airway epithelia organoid model through Air Liquid Interphase (ALI) (d0) (Fig. 5e, f). However, although Centrinone treatment during basal cell expansion in mouse cells does not impact the number of multiciliated cells (Fig. 5e, f), it causes a reduction in hybrid cilia number in terminally differentiated cells, suggesting that the hybrid cilium originates from parental centrioles (Fig. 5g, h).

### Hybrid Cilium Is a Signaling Centre and Flow Sensor

It has been previously shown that during multiciliated cells differentiation, motile cilia first generate fluid flow, then fine-tune their collective beating orientation within a cell to generate a more effective fluid flow and mucociliary transport^63,64,65^. This observation suggested the existence of a sensing mechanism converting mechanical forces into molecular signals required to fine-tune basal body orientation at the cellular level^66,67^. We reasoned that if the hybrid cilium is involved in a flow sensing mechanism, its position would be biased relative to the direction of ciliary beating. To test this hypothesis, we then developed a MATLAB image analysis script that located the position of the hybrid cilium in mature multiciliated cells relative to ciliary beating direction measured by basal body-basal foot rotational polarity (Fig. 6a). To establish whether this positional bias was dependent on fluid flow, we analyzed cells from three PCD patients with independent loss of function mutations in outer dynein arm proteins (DNAH5 and DNAH11, Sup. Table 5) critical for ciliary motility, but not basal body formation and docking. As expected, all three PCD patient cells populations exhibited normal ciliation, but impaired beating, reduced basal body rotational alignment and reduced number of aligned cells relative to healthy controls^68,69,70^(Sup. Fig. S8). Surprisingly, when the position of the hybrid cilium was measured relative to motile cilia beating direction, its position was biased toward the direction of ciliary beating in normal cells, while in airway multiciliated cells from PCD patients its position was found more centered in the cell, similarly to the position of centrosomes in cycling cells (Fig. 6b). To further confirm the biased location of the hybrid cilium, we next assessed its position relative to the cell membrane irrespective of motile cilia beating direction. Consistent with the previous analysis, in cells from PCD patients the hybrid cilium was located closer to the cell center (0.80<mean ratio<0.85), while in cells from healthy controls the hybrid cilium was found at a similar distance from the cell center and the cell membrane (mean ratio of distances to the center vs membrane = 0.97) (Fig. 6c). Altogether, our data show that the hybrid cilium position is dependent on flow direction suggesting that the hybrid cilium functions as a flow sensor.

**Figure 6.**
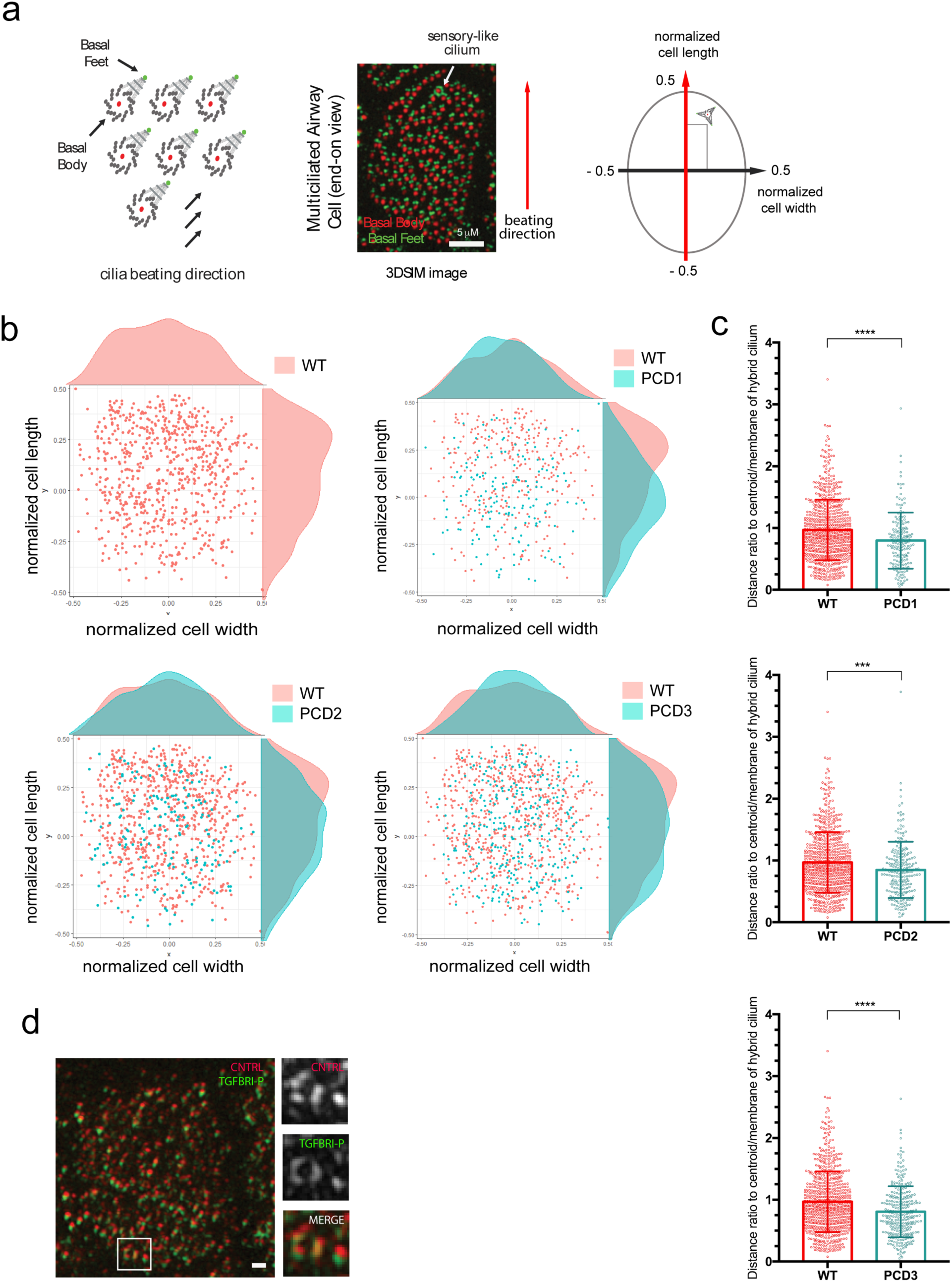
Hybrid cilium is a signaling centre whose position is dependent on flow. **(a)** Left: Cartoon depiction of the strategy for analysis of the position of the hybrid cilium. The position is calculated relative to coordinates (−0.5<x<+0.5) obtained by normalizing the cell length and width to 1. Cell length is assigned as parallel to the direction of ciliary beating measured by rotational polarity of basal bodies-basal feet pairs, cell width is assigned as perpendicular to the direction of ciliary beating. Right: MATLAB-based analysis to assess rotational polarity in multiciliated cells. **(b)** Scatterplot showing the cumulative distribution of hybrid cilia along normalized cell width and cell length in human airway multiciliated cells from healthy individuals, indicating a positional bias toward the direction of ciliary beating. Each dot represents a hybrid cilium in a cell; n=694. **(c)** Scatterplot-bar graphs showing distribution of ratio of hybrid cilium-cell centroid distance to hybrid cilium-cell membrane shortest distance of human airway multiciliated cells from healthy individuals (red, n=687) or PCD patients (PCD1-3) with immotile cilia caused by loss-of-function mutations in DNAH5 and DNAH11 (blue, n>180). Each dot represents the position of a hybrid cilium in a cell. Statistical analysis was done using Welch’s t-test. **(d)** Left: 2D projection micrograph of 3D-SIM volume of human ALI multiciliated cell treated with TGFβ1 ligand (20ng/ml; 30 min), labeled with anti-phospho-TGFBRI (green) and anti-CNTRL (red) antibodies. Right: High-magnification view of boxed area with individual channels. Scale bar represents 500 nm.

To act as a sensor, the hybrid cilium must be competent for signalling. Since basal foot protein CEP128 has been previously linked to TGFβ signalling in primary cilia^71^, we then examined whether activated TGFβ receptors were found at the hybrid cilium consistently with a sensing role. When airway multiciliated cells were labeled with an antibody recognizing phosphorylated, activated TGFβRI^72,73^ phospho-TGFβRI was found enriched at the basal bodies of airway primary and motile cilia throughout the differentiation process including at the base of the hybrid cilium (Fig. 6d). Collectively, these results suggest that the hybrid cilium is not only a fully functional motile cilium, but also a signalling antenna whose position is linked to the direction of beating.

## Discussion

### The Architecture of The Basal Foot Revealed by Super-resolution Microscopy

Here, we present the super-resolution molecular map of the basal foot and show that the basal foot has an architecture characterized by different protein regions linked by elongated coiled-coil components, which bridge different parts of the basal foot. Our data show that the basal foot in primary cilia and subdistal appendages share a conserved architecture, while basal foot in motile cilia presents a more complex structure, with arches connecting basal foot to the basal body and an overall more compact organization of region II and III relative to primary cilia. This organization most likely reflects the different mechanical requirements that the basal foot in motile cilia is subject to during ciliary beating. We initially hypothesized that the basal foot was built upon an ODF2/Cenexin molecules infrastructure. ODF2, a highly insoluble, self-interacting protein that forms a fibrillar structure is required for basal foot formation and is thought to have an indirect association with microtubules^27,74,17,33,5,4^ and recruits CEP128, TCHP, CNTRL, NIN and CEP170. However, our super-resolution map shows that ODF2 is rather a scaffold of region (II), which is then required for assembly of region III.

STORM and 3DSIM super-resolution imaging of basal foot reveals that CNTRL plays a role as a linker by connecting regions II and III along the longitudinal axis of the basal feet. In addition, in motile cilia CTRLN laterally forms two domains that are symmetrically positioned on opposite sides of the basal foot as a zipper suggesting that CTRLN provides a flexible architectural element that can be adapted to build different assemblies.

Quantitative molecular mapping directed our attention to the region of attachment of the basal boot to the basal body leading to the identification of CEP112 as a novel basal foot protein in close proximity to CEP128 and providing through BioID mapping a candidate list for future studies. Last, the basal foot map clarifies the distribution of reported basal foot proteins such as CC2D2A, TCHP and CEP19. We show that CC2D2A and TCHP are not classic basal foot proteins, a notion supported by studies of the transition zone region^75,76^. Our map also shows that CEP19 resides above the basal foot at a distance from the basal body consistent with the measurements from Kanie et al. (Sup. Fig. S5, d=334±38nm (our data); d=372.6±16.4nm^77^). This observation is supported by recent evidence suggesting that CEP19 forms a functional complex with FOP and CEP350, distal appendage proteins required for an early step of ciliogenesis^77,78,79^.

### A Novel Cilium in Airway Multiciliated Cells

Our data reveal the fate of parental centrioles in mature multiciliated cells. In airway cells, during the early stages of differentiation—before Foxj1 expression—the mother centriole templates a primary cilium that is subsequently re-adsorbed before centriole amplification^7^. After this step, its role has remained mysterious. Here we show that parental (mother) centrioles resurface to give rise to a hybrid cilium harbored among motile cilia. This hybrid cilium has features of motile and primary cilia, that is multiple basal feet as a primary cilium and a central pair apparatus and proteins required for ciliary beating as a motile cilium. Interestingly, the hybrid cilium is evolutionarily conserved in mammalian multiciliated cells, but it has not yet been identified in lower organisms such as xenopus pointing to species-specific differences.

The hybrid cilium is preferentially positioned toward the direction of ciliary beating. This biased position is reminiscent of the primary cilium at the leading edge in the wound region in vascular and bronchial smooth muscle cells, where it is thought to sense extra-cellular matrix proteins and promote cell migration^80,81^. Since effective flow generated by beating motile cilia is required to maintain the preferred position of the hybrid cilium this suggests that it senses flow either directly or indirectly. The notion of a cilium with functional attributes of motile and sensory cilia has been previously proposed in human tracheal epithelial and mouse oviduct epithelial cells^82,83^. Moreover, in lower organisms such hybrid motile-sensory cilia exist, but they have been lost during the course of evolution in higher organisms^84,85^. Consistent with a role in sensing, the parental basal body harbors phosphorylated TGFβI receptors, providing a means for signal amplification and compartmentalization around its location by possibly regulating relative intensities. TGFβ is a complex signalling pathway integrated with multiple cellular processes, whose downstream responses are cell and context dependent^86^. TGFβ signalling has been previously associated with basal foot in primary cilia^71,87^, and in multiciliated epithelia it has been shown to control motile cilia length independently from transcriptional programs responsible for multiciliogenesis^88^. It is therefore possible that the parental basal body provides a signalling hub in multiciliated cells to sense optimal/altered flow during differentiation or epithelial-to-mesenchymal transition during tissue injury or inflammation, processes linked to TGFβ signalling in airway cells^83^. Future studies are needed to address the downstream molecules and physiological effects of TGFβ signalling through the parental basal body in the airways.

## Methods

### Immortalized and Primary Cell Culture

hTERT-RPE1 cell line (source: ATC^®^ CRL-4000^TM^) was cultured in DMEM medium containing 10% FBS. HEK293 Flp-In T-Rex cells were cultured in the DMEM medium containing 10% FBS (Tetracycline-free). For ciliation, RPE-1 and HEK293 cells were serum-starved with DMEM/F-12 media for 48-72 hours. Human primary nasal airway cells from healthy volunteers and PCD patients were collected using a cytology brush by a nurse, with a protocol approved by Research Ethics Board at the Hospital for Sick Children. Airway cells were then expanded, seeded on transwells (Corning HTS Transwell-96 and −24 permeable support; 0.4 µm pore size), and differentiated for at least 21 days following Stem Cell Technologies protocols using PneumaCult-Ex and PneumaCult-ALI media. The media were supplemented with vancomycin, tobramycin, gentamicin and antibiotic-antimycotic antibiotics.

### Transfection

hTERT-RPE1 cells were transfected using Lipofectamine 3000 Kit (Invitrogen) according to manufacturer instruction. Cells were analyzed for downstream applications at 48-72 hours post transfection (hpt).

### Cloning and Plasmids

Please refer to *Supplemental materials* (Table S6) for a list of plasmids and primers used in this study.

### Antibodies

Please refer to *Supplemental materials* (Table S7) for a list of antibodies used in this study.

### Immunofluorescence

RPE-1 cells were fixed on coverslips, and human nasal and mouse tracheal multiciliated cells from ALI cultures were directly fixed on transwell filters with either methanol (20 min at −20 °C) or 4 % Paraformaldehyde (PFA; 10 min at RT). For PFA fixation, cells were subsequently reduced with 0.1% Sodium Borohydride for 7 minutes, then permeabilized with 0.2 % Triton X-100 for 25 minutes. Cells were blocked using 5% FBS-containing PBS, incubated with primary antibodies for either 1 hour (RT) or overnight (4 °C), and then secondary antibodies conjugated with Alexafluor −405, −488, −555 and −647 nm (Thermo Fisher Scientific). When appropriate, cells were stained with directly labeled primary antibodies (prepared using APEX Antibody Labelling Kit, Thermo Fisher Scientific and Mix-n-Stain Antibody Labeling Kit, Sigma-Alrich). Cells were nuclei stained with HOESCHT33342.

### Super-resolution Imaging

3DSIM data was collected using ELYRA PS.1 (Carl Zeiss Microscopy) with a Plan-Apochromat 63x or 100x/1.4 Oil immersion objective lens with an additional 1.6x optovar. An Andor iXon 885 EMCCD camera was used to acquire images with 101 nm/pixel z-stack intervals over a 5-10 µm thickness. For each image field, grid excitation patterns were collected for five phases and three rotation angles (−75°; −15°, +45°). The raw data was reconstructed and channel aligned using SIM module of ZEN Black Software (version 8.1). 2D-STORM data was collected using PALM mode in ELYRA PS.1 (Carl Zeiss Microscopy) with a Plan-Apochromat 63x or 100x/1.4 Oil immersion objective lens with an additional 1.6x optovar. An Andor iXon 885 EMCCD camera was used to acquire images using TIRF mode. Lasers of wavelength 647 nm and 405 nm (if necessary) were used to activate the fluorophore. Raw data was reconstructed using PALM module of Zen Black Software (version 8.1), with the account for overlapping molecules. Reconstructed data was further processed for drift correction and binning using home-written MATLAB script (can be accessed via the following link: https://drive.google.com/open?id=11fuWn7kmZ-loCn79CKChJI5FeMme0fDU).

### Transmission Electron Microscopy (TEM)

ALI filters of fully differentiated human nasal multiciliated cells were fixed in 2% glutaraldehyde in 0.1M sodium cacodylate buffer. Samples were rinsed in 0.1M sodium cacodylate buffer with 0.2M sucrose, post-fixed in 1% OsO_4_ in 0.1M sodium cacodylate buffer, dehydrated in a graded ethanol series (70%, 90%, 3X 100%), infiltrated with propylene oxide, and embedded in Quetol-Spurr resin. Serial sections (90 nm-thickness) were cut on a Leica Ultracut ultramicrotome, stained with uranyl acetate and lead citrate, and imaged in a FEI Tecnai 20 TEM.

### Focused Ion Beam Scanning Electron Microscopy (FIB-SEM)

ALI filters of fully differentiated human nasal multiciliated cells were fixed in 2.5% glutaraldehyde and 0.05% malachite green oxalate in 0.1M sodium cacodylate buffer, rinsed in 0.1M sodium cacodylate buffer, post-fixed in 0.8% potassium ferrocyanide and 1% OsO_4_ in 0.1M sodium cacodylate buffer. The samples were treated with 1% tannic acid, stained with 0.5% uranyl acetate, followed by dehydration in a graded acetone series (25%, 50%, 75%, 95% and 100%), and embedded in resin. Resin formulation: 18.2% Araldite M (Sigma-Aldrich), 22.7% Epon 812 (Sigma-Aldrich), 54.5% Hardener DDSA (Sigma-Aldrich) and 4.5% DMP-30 (Sigma-Aldrich). FIB-SEM imaging for Sup. Movie 1,2 was performed as described below. Sample blocks for analysis by FIB-SEM were trimmed and mounted on a 45° pre-titled SEM stub and coated with a 4-nm layer of Pt to enhance electrical conductivity. Milling of serial sections and imaging of block face after each *Z*-slice was carried out with the FEI Helios Nanolab 660 DualBeam using Auto Slice & View G3 ver 1.5.3 software (FEI Company, Hillsboro, OR USA). A block was first imaged to determine the orientation relationship between the block face of ion and electron beams. A protective carbon layer 50 μm long, 8 μm wide and 2 μm thick was deposited on the surface of the region of interest to protect the resin volume from ion beam damage and correct for stage and/or specimen drift, i.e., perpendicular to the image face of the volume to be milled. Trenches on both sides of the region of interest were created to minimize re-deposition during automated milling and imaging. Imaging fiducials were generated for both ion and electron beam imaging and were used to dynamically correct for drift in the x-and y-directions by applying appropriate SEM beam shifts. Ion beam milling was performed at an accelerating voltage 30 kV and beam current of 9.3 nA, stage tilt of 9°, and working distance of 4 mm. With each milling step, 10 nm thickness of the material was removed. Each newly milled block face was imaged with the through-the-lens detector for backscattered electrons (TLD-BSE) at an accelerating voltage of 2 kV, beam current of 0.4 nA, stage tilt of 47°, and working distance of 3 mm. The pixel resolution was 10.3 nm with a dwell time of 30 μs. Pixel dimensions of the recorded image were 1536 x 1024 pixels. Seven hundred and forty-three images were collected and the images contrast inversed. Visualization and direct 3-D volume rendering of the acquired dataset was performed with Amira 6.0.1 (FEI Company, Hillsboro, OR USA). FIB-SEM imaging for Sup. Movie 3 was performed as describe previously^89^.

#### Western Blot

Total cell lysates were collected using RIPA lysis buffer (Pierce, Thermo Fisher Scientific) freshly added with protease inhibitor (Roche, Sigma-Aldrich). Lysates were loaded on 4-12 or 8% Bis-Tris Plus gels. Proteins were transferred to a nitrocellulose membrane and blocked using 5% Skim-milk in TBST. Protein blots were sequentially incubated with primary and HRP-conjugated secondary antibodies diluted in 5% BSA in TBST. Blots were developed using the Novex ECL Chemiluminescent Substrate Kit (Invitrogen).

### Bio-ID Assay

HEK293 Flp-In T-Rex cells were first co-transfected with the pcDNA5/FRT/TO CEP128-FLAG-BirA* plasmid and Flp Recombinase Expression plasmid pOG44 (1:20 ratio) and selected for stable expression with Hygromycin B and Blasticidin. Stable CEP128-FLAG-BirA* HEK293 Flp-In T-Rex cell line was induced for BirA expression and biotinylated for 24 hrs with 1 µg/ml tetracycline and 50 µM biotin. For ciliation experiments, cells were serum-starved to for 72 hrs. Cells were then collected and processed for Bio-ID and FLAG Immunoprecipitation (IP) experiments as described previously^90^.

### MTEC and ependymal cell experiments

Mouse tracheal epithelia cell (MTEC) cultures were established as previously described^91,92^. Briefly, C57BL/6 mice were sacrificed at 2–4 months of age, trachea were excised, opened longitudinally to expose the lumen, and placed in 1.5 mg/mL Pronase E in DMEM/F12 medium (Life Technologies) at 4°C overnight. Tracheal epithelial cells were dislodged by gentle agitation and collected in DMEM/F12 with 10% FBS. After centrifugation, cells were treated with 0.5 mg/mL DNase I for 5 min on ice and centrifuged at 4°C for 10 min at 400 g. Cells were resuspended in DMEM/F12 with 10% FBS and plated in a tissue culture dish for 5 h at 37°C with 5% CO_2_ to adhere contaminating fibroblasts. Non-adhered cells were then collected, concentrated by centrifugation, resuspended in an appropriate volume of mTEC-Plus medium^92^, and seeded onto Transwell-Clear permeable filter supports (Corning).

To eliminate parental centrioles, cells were incubated in the presence of 1 μM Centrinone A^62^ for 6 days. Air-liquid interface (ALI) was established 2 days after cells reached confluence by feeding mTEC-Serum-Free medium^92^ only in the lower chamber. Cells were cultured at 37°C with 5% CO_2_, and media replaced every 2 d, and fixed on the indicated days. All chemicals were obtained from Sigma Aldrich unless otherwise indicated. Media were supplemented with 100 U/mL penicillin, 100 mg/mL streptomycin, and 0.25 mg/mL Fungizone (all obtained from Life Technologies).

For ependymal cell culturing, all animal studies were performed in accordance with the guidelines of the European Community and French Ministry of Agriculture and were approved by the Ethic comity Charles Darwin (C2EA-05) and “Direction départementale de la protection des populations de Paris”, (Approval number Ce5/2012/107; APAFiS #9343). The mouse strain, Cen2-GFP (CB6-Tg (CAG-EGFP/CETN2)3-4Jgg/J, The Jackson Laboratory), has already been described^61^. For *in vivo* analysis, animals used were homozygous for the Cen2-GFP. Lateral walls of the lateral brain ventricles were dissected as previously explained^93^. The tissue was treated with 0.1% triton in BRB80 (80 mM K-Pipes pH6.8; 1 mM MgCl2; 1 mM Na-EGTA) for 1 min prior to fixation and fixed in methanol at −20°C for 10 min. Saturation and antibody incubations were performed in PBS containing 10% FBS and 0.1% triton. Primary antibodies (CNTRL (monoclonal mouse from Santa Cruz) and CEP164) were incubated overnight (4°C). Secondary antibodies conjugated with Alexa Fluor −555 and −647 were incubated for 1h.

For *in vitro* pulse-chase experiments, cultures were performed as previously described. Transfection of ependymal cell progenitors was performed at 80% of confluency during the proliferation phase with a CMV-TagRFP-Cen1 plasmid (gift from Xavier Morin, ENS, Paris), which codes for human centrin 1 fused to TagRFP under the control of a CMV promoter, using jetPRIME Polyplus kit. Cells (in 25cm^3^ flask) were transfected with a mix of 0.75µg of DNA, 300µL of jetPRIME Buffer and 1.5µL of jetPRIME transfection reagent in 3 mL of fresh complete medium (DMEM-Glutamax (Invitrogen) containing 10% FBS and 1% Penicillin/Streptomycin). After 4 hours at 37 °C in 5% CO2 incubator, the medium was renewed. One day after proliferation, cells were shaken at 250 rpm overnight. Cells were plated on coverslips or Labtek chambers slides coated with L-Polylysin (40 μg/ml in pure water) at a density of 0.75 × 10^4^ cells per µl in 20 or 60 µl drops. The medium was then replaced by serum-free DMEM-Glutamax-I 1% P/S, to trigger ependymal differentiation in vitro (DIV0). Cells were either fixed with Paraformaldehyde (4% in PBS) for 10min or used for live imaging. Fixed cells were examined with an upright epifluorescence microscope (Zeiss Axio Observer.Z1) equipped with Apochromat X63 (NA 1.4) or X100 (NA 1.4) oil-immersion objectives and a Zeiss Apotome with an H/D grid. Images were acquired using Zen software with 230-nm z-steps and analyzed with image-J.

For live imaging, differentiating ependymal cells with two bright RFP-tagged centrosomal centrioles were selected and filmed using an inverted spinning disk Nikon Ti PFS microscope equipped with an oil-immersion X100 (NA 1.4) objective, an Evolve EMCCD Camera (Photometrics), dpss lasers (491 nm, 561 nm), a motorized scanning deck and an incubation chamber (37 °C; 5% CO2; 80% humidity). Laser intensities and image capture times were respectively set to 20%, 50 ms for 488nm and 25%, 100 ms for 561nm. Images were acquired with Metamorph software at 60 minutes time interval for 24 hours. Image stacks were recorded with a z-distance of 0.7 mm. Four dimensional (x, y, z, t) time-lapse images were analyzed with Image J.

#### Semi-quantitative RT-PCR

RNA was purified from 1.5 x 10^5^ cells on coverslips using the RNeasy Micro Kit (QIAGEN, 74000). Retrotranscription was performed using SuperScript III First-Strand Synthesis System for RT-PCR (Invitrogen, 18080-051). PCR was performed on cDNA using the primers 5’-AGAAGAACGGCATCAAGGTG-3’ and 5’-GAACTCCAGCAGGACCATGT-3’ for EGFP 5’-AACACCGAGATGCTGTACCC-3’ and 5’-ACGTAGGTCTCTTTGTCGGC-3’ for tagRFP and 5’-ACCCCACCGTGTTCTTCGAC-3’ and 5’-CATTTGCCATGGACAAGATG-3’ for cyclophilin. Images of the gels were then analysed on ImageJ. The ratio between EGFP or tagRFP and cyclophilin band intensity were calculated. Quantifications of 3 independent experiments were pooled and plotted.

### Rotational polarity assessment and positional analysis of hybrid cilium

Custom written MATLAB script was used to determine the position of the hybrid cilia in multiciliated cells relative to cilia beating direction (can be accessed via the following link: https://drive.google.com/open?id=182KAccJf6YC69WbovKgTwtg62Y5DTadA). First, intensity thresholds for all channels were chosen for and binary images were generated to identify individual basal body and basal foot objects. Individual cells were outlined via manual cell border drawing. Basal body-basal foot pairs were identified based on the pairwise nearest neighbor search with a distance threshold of ∼600 nm. The direction of a single cilium was defined as from the weighted center of the basal body object to that of the paired basal foot. All cilia directions in one cell were determined and the mean direction was regarded as the direction of beating in a cell. The cilia beating angles obtained were transformed into a two-dimensional unit vector: 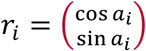. The resultant vector was the average of all the unit vectors in a cell: 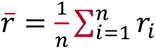. The resultant vector length r was defined as the norm of the resultant vector: 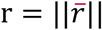. The circular standard deviation was defined as: 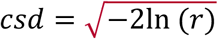. All directions in a single cell were also subject to Rayleigh’s test for uniformity distribution. The p-value is calculated as: 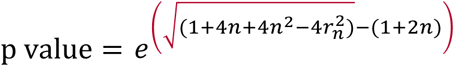; *r*_*n*_= *r* × *n*. A p-value < 0.05 indicated that the cilia in the cell are significantly aligned. Aligned vector length was defined to describe the cilia alignment level in a cell with values ranging from 1 to 0, with 1 indicating 100% alignment and 0 indicating no alignment. The mean beating direction of all cilia were defined as the cilia beating direction. The hybrid cilia position relative to the cilia beating direction was measured using the same basal foot and basal body markers in cells whose size is normalized to [-0.5; 0.5] both along the cilia beating direction (regarded as cell length) and the direction perpendicular to it (regarded as cell width).

### Statistical Analysis

Data was analyzed in Microsoft Excel and Prism software. Statistical tests, sample sizes and number of replicates were specified in figure legends. Differences were regarded as significant if p < 0.05, unless otherwise stated.

## Supporting information

## Acknowledgements

This project is funded by CIHR program grant # 391917 to VM and SD; National Heart, Lung and Blood Institute (R01-HL128370) and National Institutes of Diabetes and Digestive and Kidney Diseases (R01-DK108005) to MRM; Z.L. was supported by the SickKids Restracomp Fellowship; The authors acknowledge PCD patients and volunteers for providing nasal cells for this study, Julie Avolio for help with nasal cell scraping. Jia Zhou, Cindy Fang, Alexandra Albulescu and Jasmine Kang assisted in data analysis. Douglas Holmyard (EM facility, The Hospital for Sick Children) prepared TEM and FIB-SEM samples and helped set up EM imaging. McGill EM facility for FIB-SEM acquisition. We thank Profs Bornens, Pelletier, Cheeseman, Kyung Lee, Elsasser, Avidor-Reiss for generously sharing antibodies and plasmids.

## Author Contributions

Q.P.H.N. designed and conducted experiments, collected and analyzed the data, wrote the manuscript. Z.L. wrote MATLAB scripts and did STORM experiments. L.Z., H.O., W.F. and T.M. helped with airway multiciliated cells culturing system; RN performed the mouse tracheal epithelia cell experiments; MM and VM conceived and designed the MTEC-centrinone experiments; N.D., M.A and A.M. conceived and performed the ependymal cells pulse chase experiments; E.C., E.L. and B.R. performed BioID experiments; S.D. provided PCD patients cells, K.C. helped with sample preparation for FIB-SEM data collection and acquired data; V.M. conceived the project, designed experiments, collected data, analyzed data and wrote the manuscript.

## Supplemental Tables

**Table S1:**
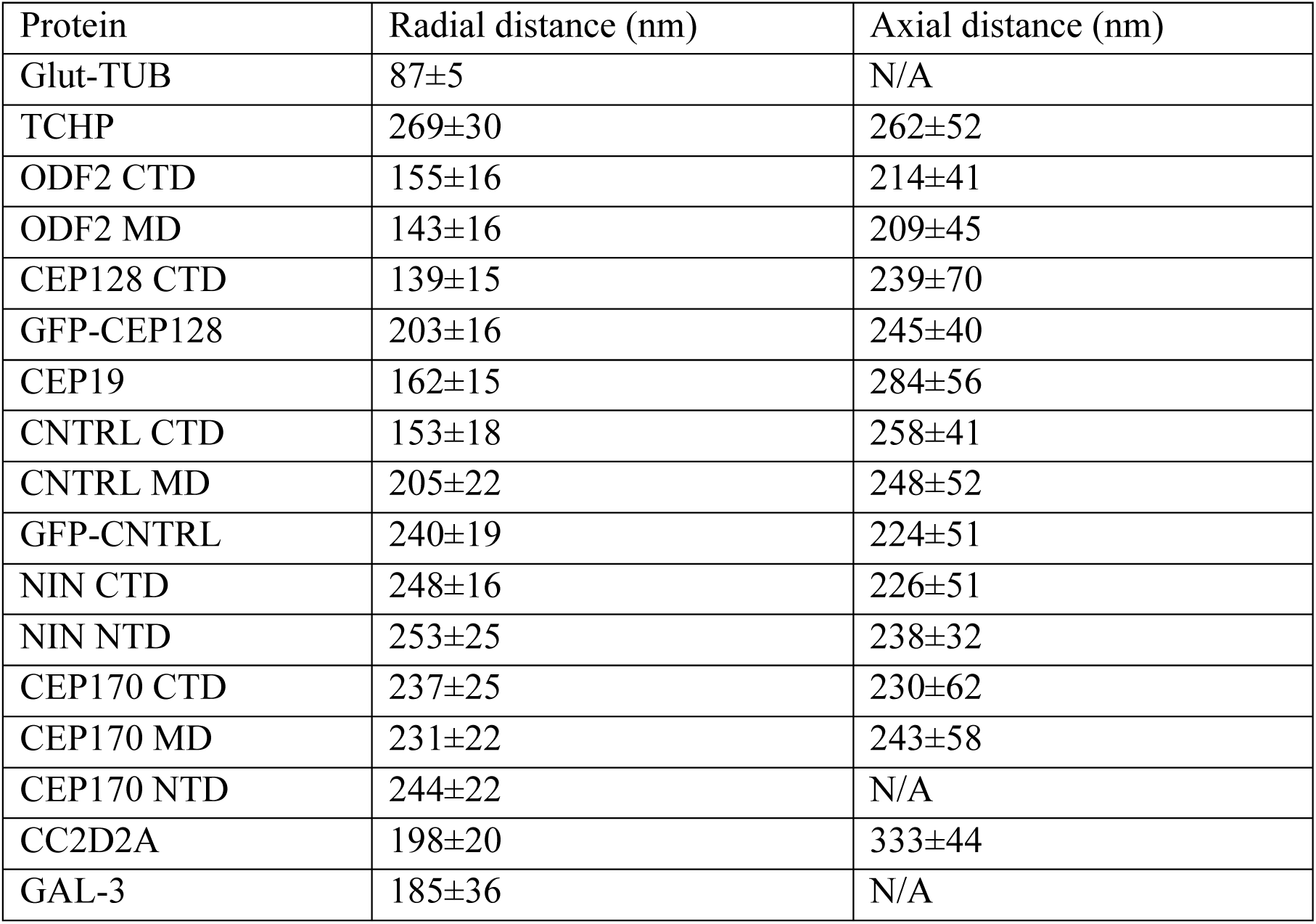
Radial and axial distance measurements of basal foot proteins in primary cilia (n=40)

**Table S2:** Radial distance measurements of GFP-NIN constructs in ciliated RPE cells (n>23)

| NIN constructs | radial distance (nm) |
| --- | --- |
| GFP-NIN | 245±25 |
| GFP-aa197 | 218±32 |
| GFP-aa764 | 213±32 |
| GFP-aa1640 | 213±36 |
| GFP-aa1647 | 201±32 |

**Table S3:** Radial distance measurements of subdistal appendage proteins in cycling cells (n=40)

| protein | radial distance (nm) |
| --- | --- |
| ODF2 CTD | 137±16 |
| CEP128 CTD | 144±16 |
| CNTRL CTD | 171±16 |
| NIN CTD | 245±18 |
| NIN NTD | 243±25 |
| CEP170 CTD | 231±20 |
| CEP170 MD | 249±21 |

**Table S4:** Radial distance measurements of basal foot proteins in motile cilia (n=80)

| protein | radius distance (nm) |
| --- | --- |
| ODF2 CTD | 158±16 |
| ODF2 MD | 147±5 |
| CEP128 CTD | 162±25 |
| CEP19 | 177±22 |
| CNTRL CTD | 180±23 |
| CNTRL MD | 200±224 |
| NIN CTD | 188±29 |
| NIN NTD | 177±32 |
| CEP170 CTD | 191±30 |
| CEP170 NTD | 194±33 |
| NEDD1 | 197±41 |
| Gamma-tub | 206±33 |

**Table S5:** PCD patients genotype

| Name | Gene | Mutation(s) |
| --- | --- | --- |
| PCD1 | *DNAH5* | c.[1432>T]; c[11571-iG>A] |
| PCD2 | *DNAH11* | p.[(Cys4286*)];[(Ile4122Ser)] |
| PCD3 | *DNHA5* | Hom p.[(Phe634Serfs*2)] |

**Table S6:**
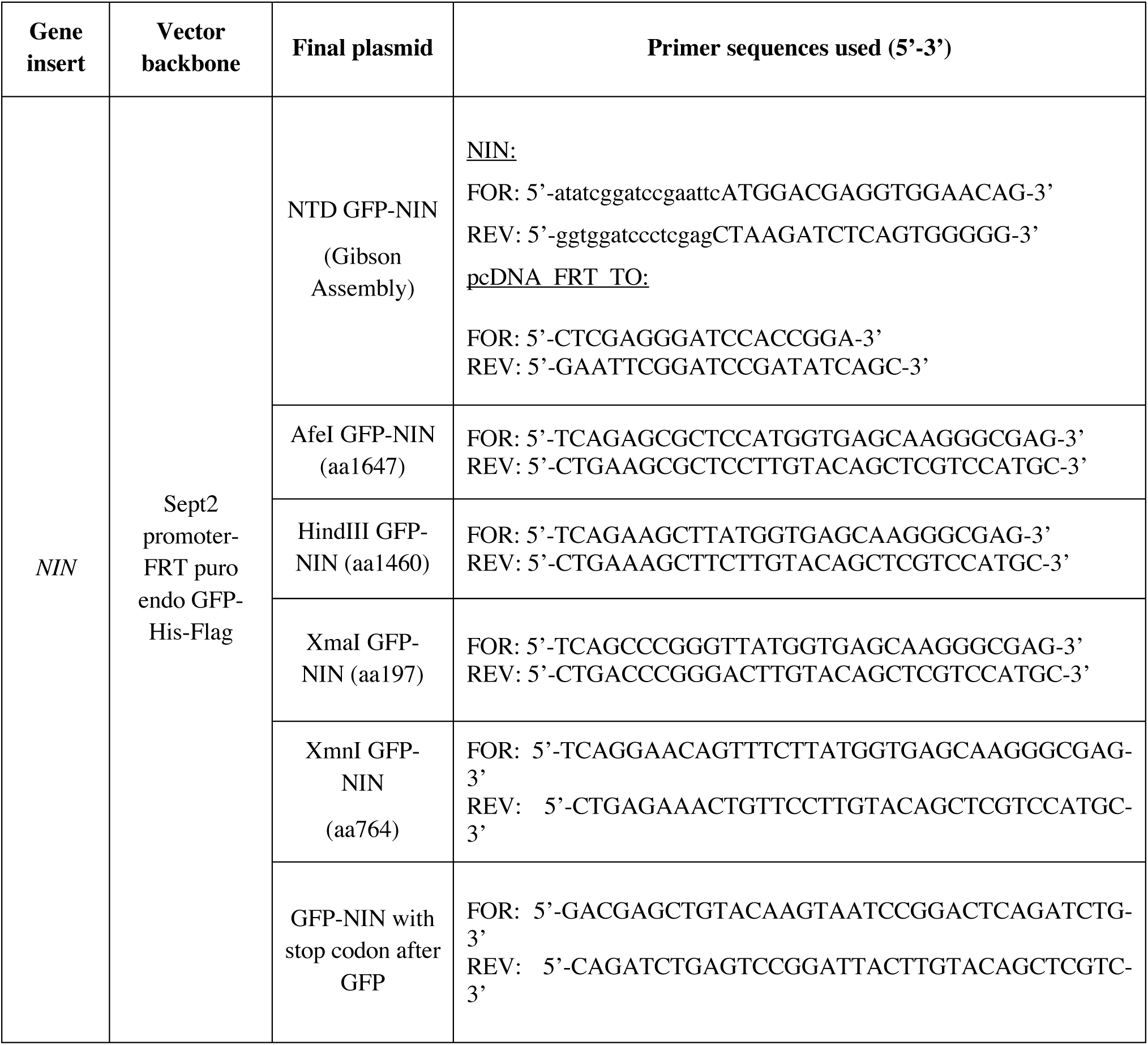
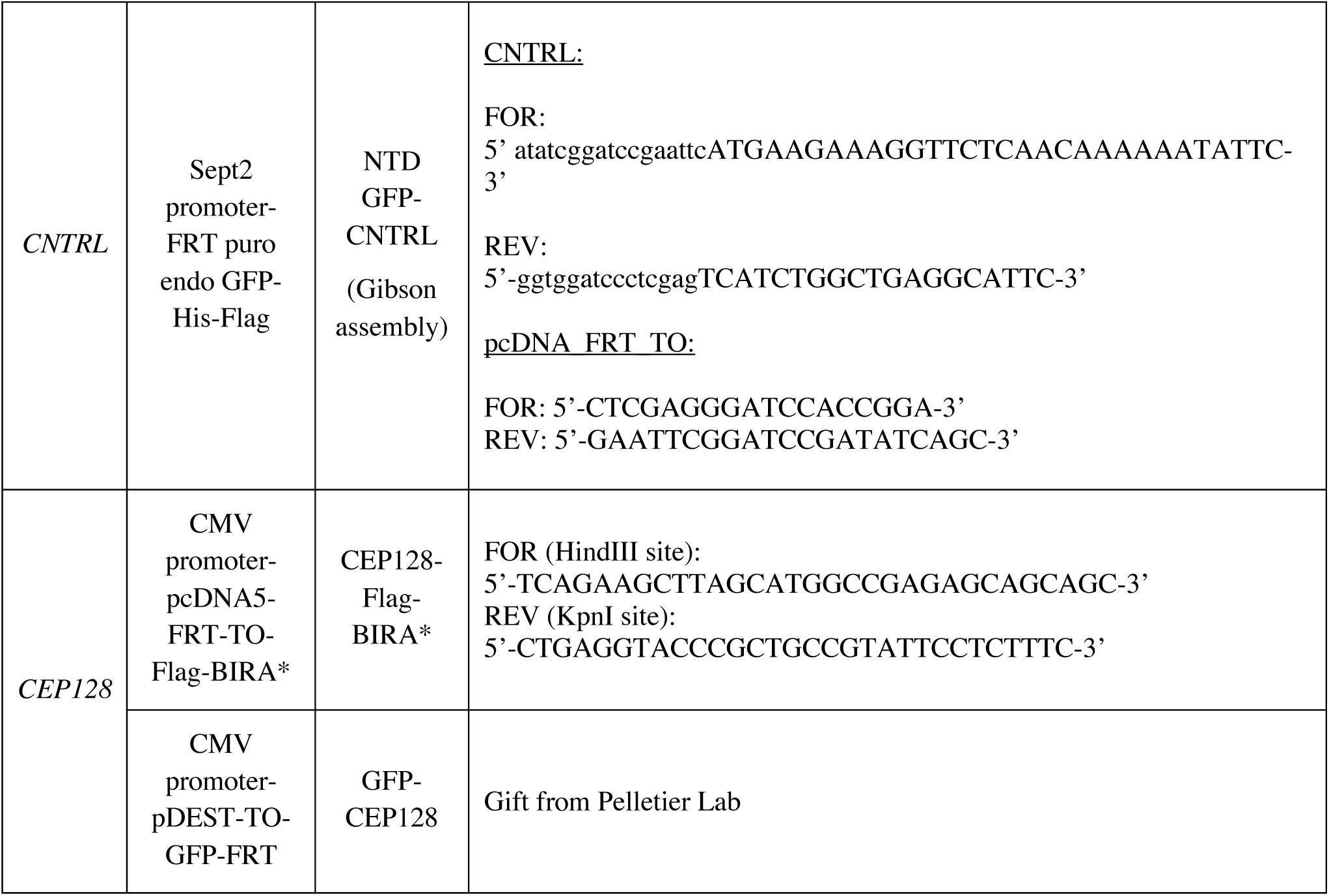
List of plasmid and primers used in this study

**Table S7:** List of antibodies used in this study Note: CTD-recognizing antibodies were used at default in case of multiple antibodies for the same protein, unless mentioned otherwise.

| Gene | Domain recognized | Source | Catalog # | Species | Dilution | Fixation method |
| --- | --- | --- | --- | --- | --- | --- |
| *CEP128* | CTD (aa 1044-1094) | Abcam | Ab11879 | rabbit | 1:50 | methanol |
| *ODF2* | CTD (aa 750-C-terminus) | Abcam | Ab43840 | rabbit | 1:500 | methanol |
|  | NTD<br>aa 250–632<br>(hCenexin1) | Kyung Lee Lab | - | rabbit | 1:50 | methanol |
|  | aa 255–618<br>(hOdf2) |  |  |  |  |  |
| *CEP19* | aa 1-163 | Proteintech | 26036-1-AP | rabbit | 1:50 | methanol |
| *CNTRL* | CTD (aa 2026-2325) (for STORM experiment) | Santa Cruz | sc-135020 | rabbit | 1:50 | methanol |
|  | CTD (aa 2187-2291) | Atlas Antibodies | HPA020468 | rabbit | 1:50 | methanol or PFA |
|  | CTD (aa 2026-2325) | Santa Cruz | sc365521 | mouse | 1:50 | methanol |
|  | MD (aa 1063-1168) | Atlas Antibodies | HPA051583 | rabbit | 1:25 | methanol |
| *NIN* | CTD (aa 1870-2020) | Proteintech | 13007-1-AP | rabbit | 1:500 | methanol |
|  | NTD (L77) | Michael Bornens Lab | - | rabbit | 1:10,000 | methanol |
|  | NTD (in SDA measurements) | Santa Cruz | sc-5014 (Y-16) | goat | 1:50 | methanol |
| *CEP170* | CTD (aa 1534-C-terminus) | Abcam | Ab7250 | rabbit | 1:200 | methanol |
|  | MD (aa 1010-1200) | Iain Cheeseman Lab | - | rabbit | 1:1000 | methanol |
|  | NTD | Thermo Fisher | 41-3200 | mouse | 1:200 | methanol |
| *CCDC120* | aa 150-250 | Atlas Antibodies | HPA000561 | rabbit | 1:50 | methanol |
| *CCDC68* | aa 165-247 | Atlas Antibodies | HPA048197 | rabbit | 1:50 | methanol |
| *GAL3* | - | Elsasser Lab | - | rabbit | 1:100 | methanol |
| *CC2D2A* | - | Proteintech | 22293-1-AP | rabbit | 1:50 | methanol |
| *CEP112* | CTD (aa 607-955) | Proteintech | 24928-1-AP | rabbit | 1:100 | methanol |
|  | MD (aa361-427) | Atlas antibodies | HPA024481 | rabbit | 1:50 | methanol or PFA |
| *POC1B* | aa 321-350 | Invitrogen | PA5-24495 | rabbit | 1:50 | methanol |
|  | - | Tomer Avidor-Reiss Lab | - | rat | 1:50 | methanol |
| *Polyglutamylation Modification* | - | Adipogen | AG-20B-0020-C1 (GT335) | mouse | 1:1000 | methanol |
| *TUBA4A (α-tubulin)* | - | Sigma-Aldrich | T9026 (DM1A) | mouse | 1:500 | methanol or PFA |
| *TUBA4A (α-tubulin- FITC conjugated)* | - | Sigma-Aldrich | F2168 (DM1A) | mouse | 1:500 | methanol or PFA |
| *SAS6* | - | Santa Cruz | sc-82360 | mouse | 1:200 | methanol |
| *TCHP* | aa 261-335 | Atlas Antibodies | HPA038638 | rabbit | 1:50 | methanol |
| *GFP* | full-length | Abcam | Ab13970 | chicken | 1:2000 | methanol |
| *Turbo GFP* | full-length | Thermo Fisher | PA5-22688 | rabbit | 1:500 | PFA |
| *NEDD1* |  | Abcam | 57336 | mouse | 1:500 | methanol |
| *γ-tubulin* |  | Sigma | T6557 | mouse | 1:500 | methanol |
| *RPGRIP1L* |  | Proteintech | 55160-1-AP | rabbit | 1:50 | methanol |
| *RSPH4A* |  | Atlas Antibodies | HPA031196 | rabbit | 1:50 | methanol |
| *GAS8* |  | Atlas Antibodies | HPA041311 | rabbit | 1:50 | methanol |
| *CEP120* |  | Moe Mahjoub Lab | Mahjoub et al. JCB 2010 | rabbit | 1:10000 | methanol |
| *CEP164* | - | Novus Biologicals | 45330002 | rabbit | 1:500 | methanol |
| *acetylated α-tubulin* |  | Sigma | T7451 | mouse | 1:50 | methanol |
| *ZO-1* |  | Santa Cruz | sc-33725 | rat | 1:1000 | methanol |
| *phospho-TGFBeta RI (S165)* | aa 131-180 | Abcam | ab112095 | rabbit | 1:50 | PFA |

