## Supplementary material for "Super-resolution Molecular Map of Basal Foot Reveals Novel Cilium in Airway Multiciliated Cells"

**Supplemental Figures**

### Figure S1: Localization of basal foot proteins in primary cilia

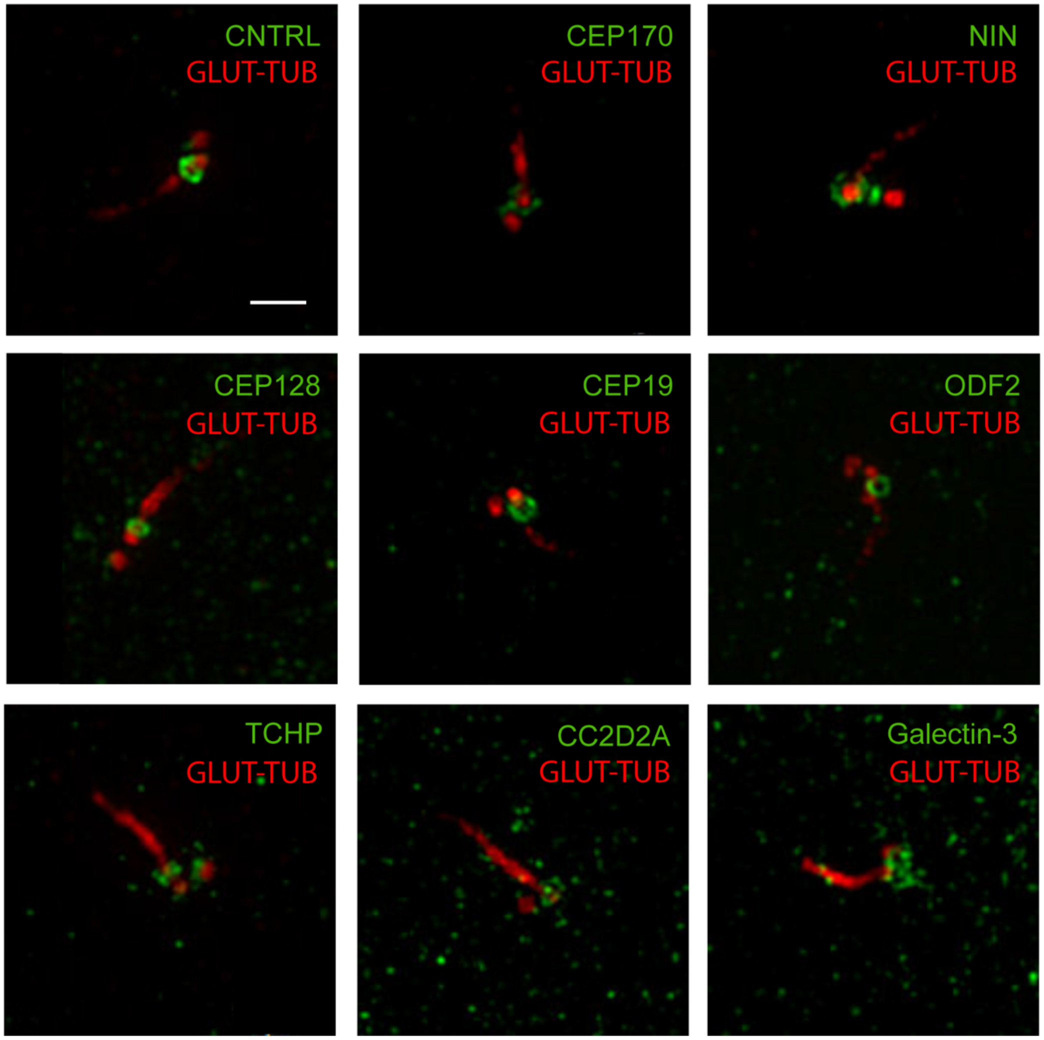


**Figure S1. Localization of basal foot proteins in primary cilia**

2D projection micrographs of 3DSIM volumes of ciliated RPE-1 cells labeled with anti-basal foot proteins (green) and anti-glutamylated tubulin (red) antibodies. Scale bar represents 1 μm.

### Figure S2. Ninein has an elongated looping distribution that connects region 2 and 3

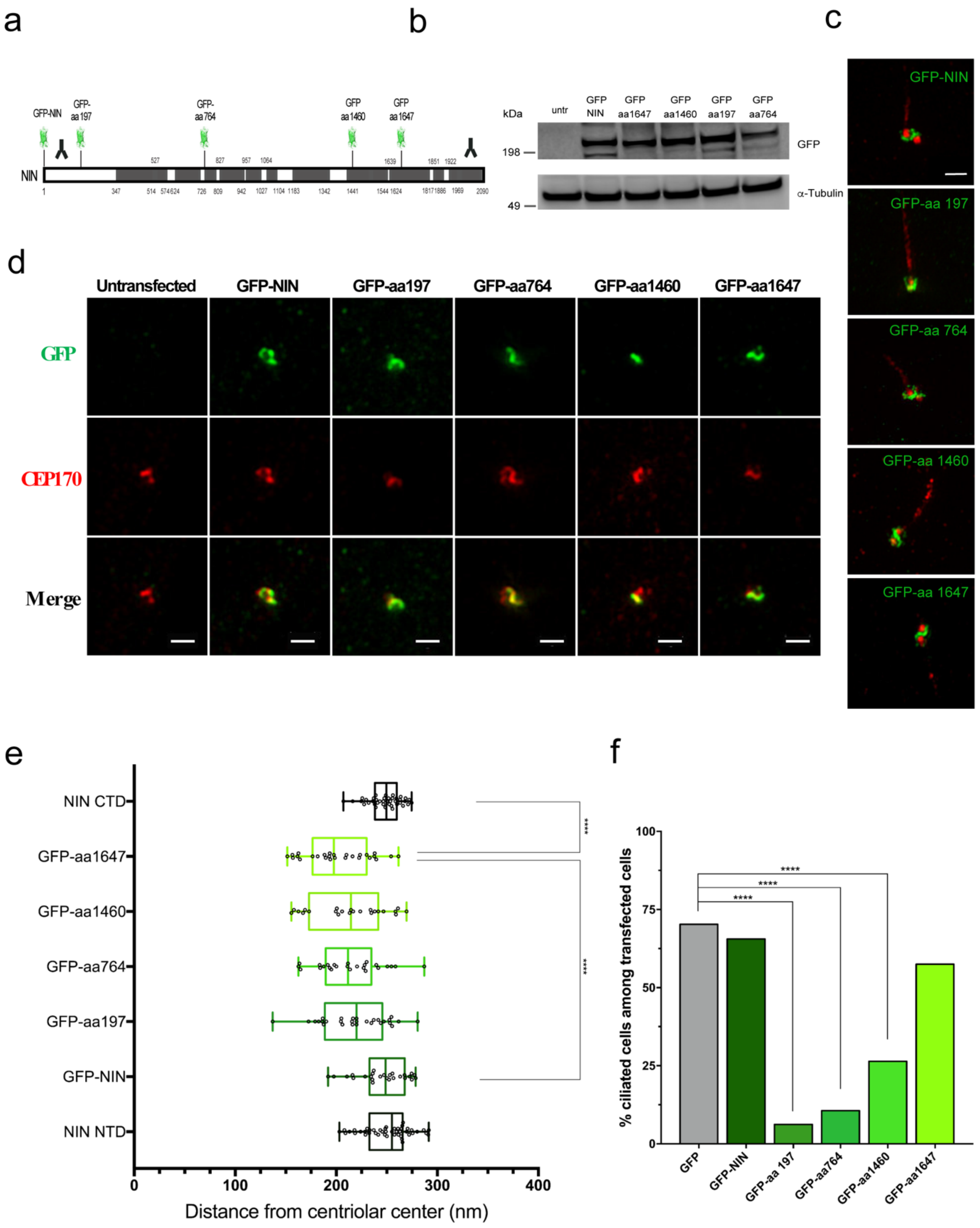

**Figure S2. Ninein has an elongated looping distribution that connects regions 2 and 3**

**(a)** Linear map representing protein polypeptide sequence of NIN with regions recognized by antibodies and positions of GFP-insertion sites. **(b)** Western blot of RPE-1 lysates from control untransfected cells or from transfected cells with different Ninein-GFP constructs for 72h. Membrane were probed with anti-GFP (Ninein: 248kDa) and anti-alpha tubulin antibodies. **(c)** 2D projection micrographs of 3DSIM volumes of RPE-1 cells expressing different Ninein-GFP constructs (72 hpt), labelled with anti-GFP (green) and anti-glutamylated tubulin (red) antibodies. Scale bar represents 1 μm. **(d)** 2D projection micrographs of 3DSIM volumes of RPE-1 cells transfected with different GFP-NIN constructs, labelled with anti-GFP (green) and anti-CEP170 (red) antibodies. Note the normal recruitment of CEP170 upon transfection of different NIN-GFP constructs. Scale bar represents 1 μm. **(e)** Box plot of radial distance of different Ninein-GFP constructs in RPE-1 cells, relative to C- and N- terminally labeled Ninein (n>23). Statistical analysis was done using one-way ANOVA and Tukey's multiple comparison test. **(f)** Bar graph showing percentage of ciliated RPE-1 cells expressing different GFP-NIN constructs over total transfected population (n>1000). GFP-expressing cells were used as a control. Statistical analysis was conducted using Fisher's exact test.

### Figure S3: The conserved architecture of subdistal appendage and basal foot of primary cilia

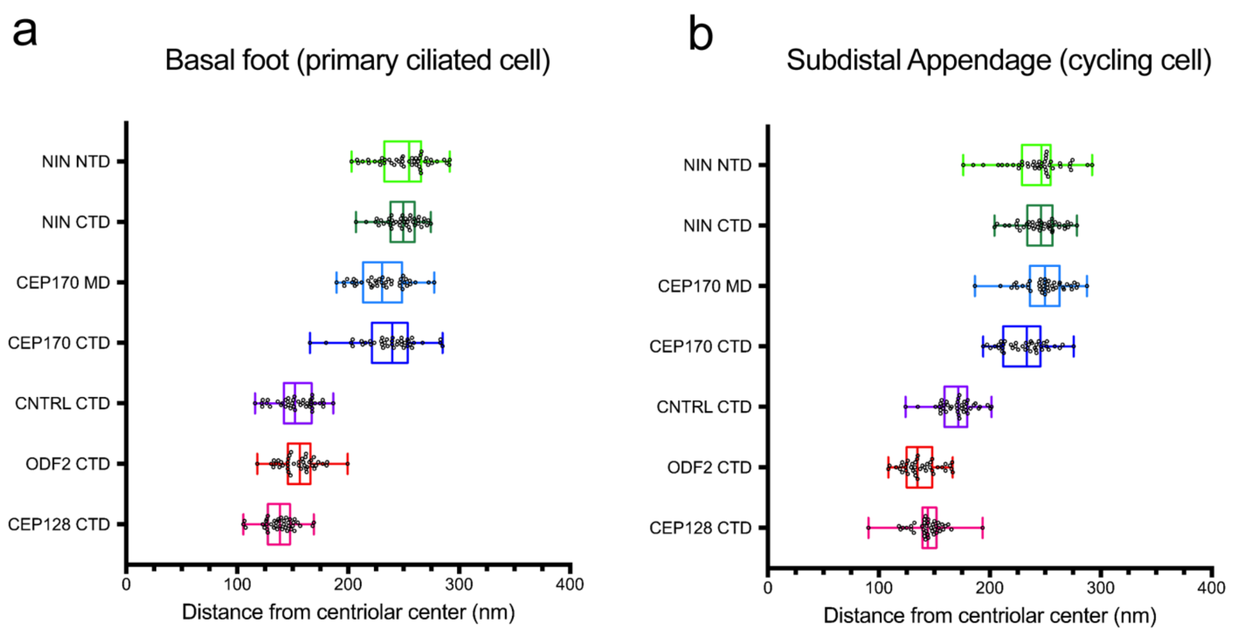


**Figure S3. The conserved architecture between subdistal appendage and basal foot of primary cilia**

Box plot of radial distributions of subdistal appendage/basal foot proteins in (a) ciliated, and (b) cycling RPE-1 cells (n=40).

### Figure S4. CEP112 antibody validation by Western blot

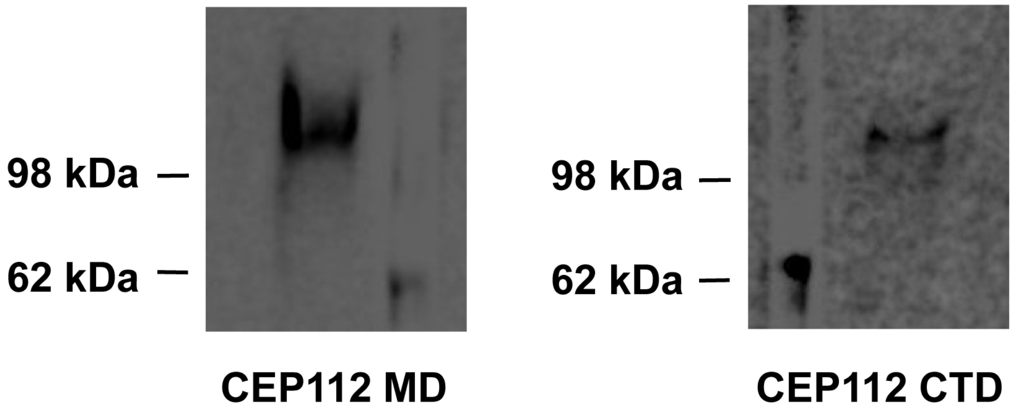


**Figure S4. Anti-Cep112 antibody validation**

Western blot of RPE-1 lysates from cells serum-starved for 72 hours to induce ciliation. Membrane were probed with anti-Cep112 antibodies raised against the MD (left and CTD (right) of the protein.

### Figure S5: CEP112, CEP19 and TCHP localization in motile cilia

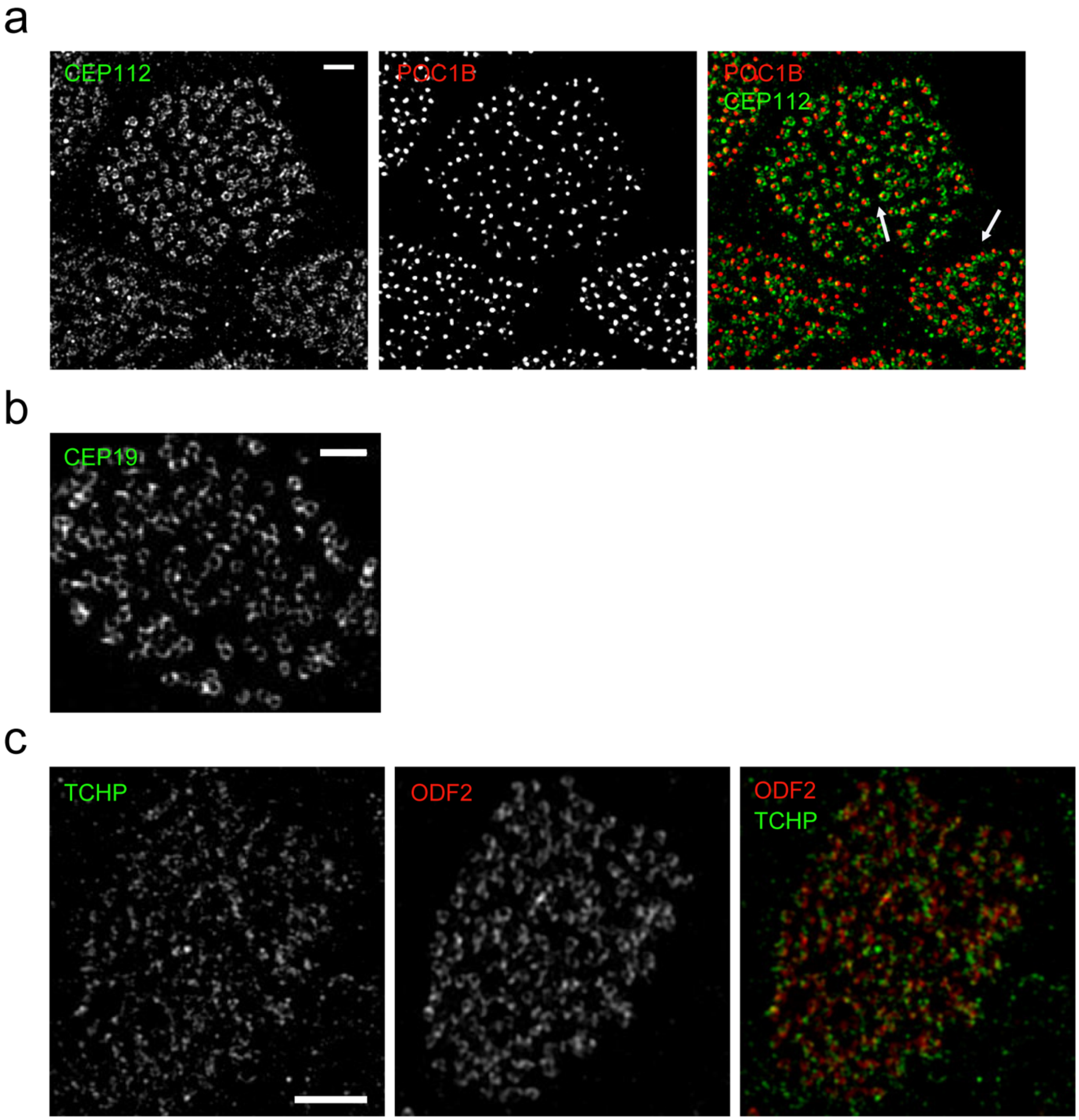

**Figure S5. CEP112, CEP19 and TCHP localization in motile cilia**

**a)** 2D projection micrograph of 3DSIM volume of human airway multiciliated cells, labeled with anti-CEP112 CTD (green), anti-poc1b (red) antibodies. Scale bar represents 1 μm. **b)** 2D projection micrograph of 3DSIM volume of human airway multiciliated cells, labeled with anti-CEP19 antibodies. Scale bar represents 1 μm. **c)** 2D projection micrograph of 3DSIM volume of human airway multiciliated cells, labeled with anti-TCHP (green) and anti-ODF2 (red). Scale bar represents 2 μm.

### Figure S6. CEP128 and ODF2 localization at hybrid cilia in human airway multiciliated cells

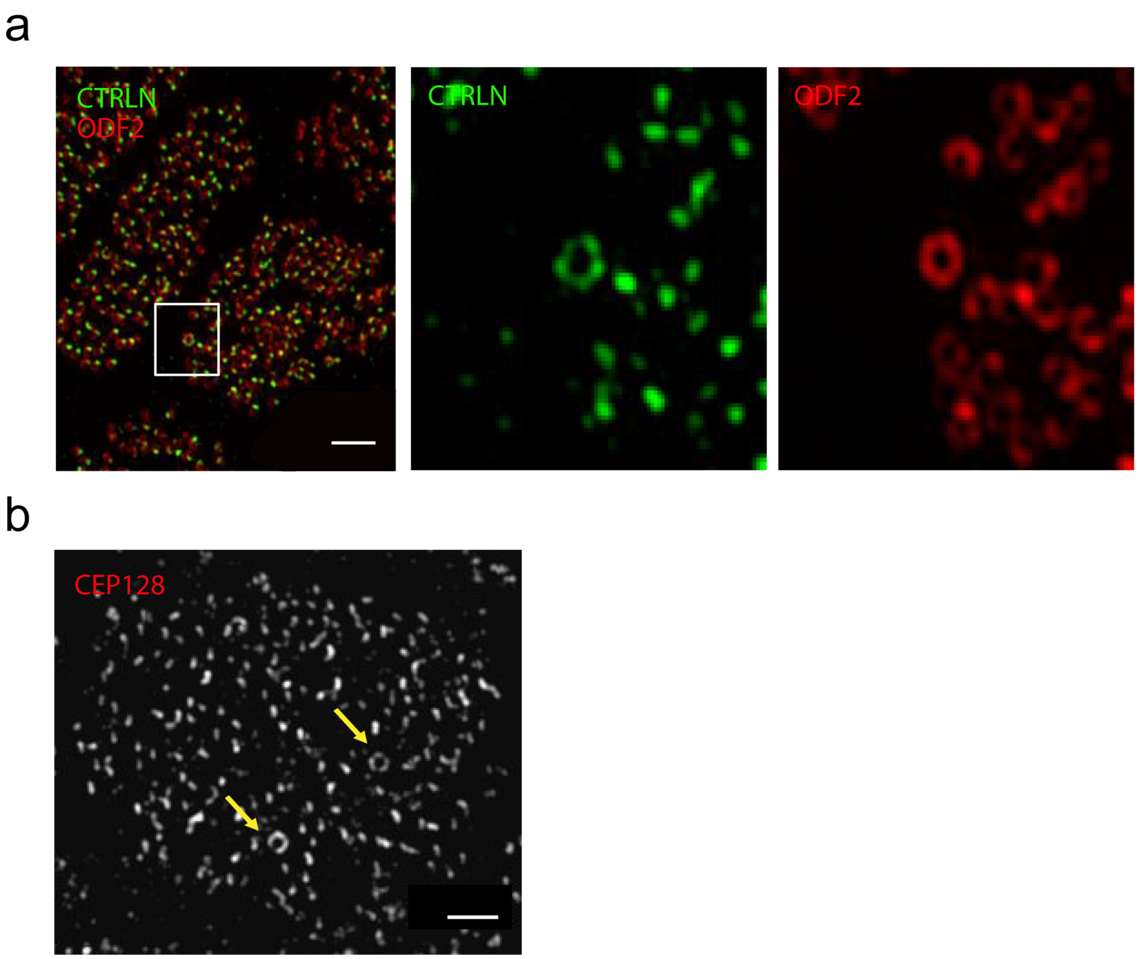


**Figure S6. ODF2 and CEP128 localization at hybrid cilia in human airway multiciliated cells**

**a)** Left: 2D projection micrograph of 3DSIM volume of human airway multiciliated cells. Right: high-magnification view of the boxed area with individual channels (middle and right), labeled with anti-CNTRL (green) and anti-ODF2 (red) antibodies, showing a ring-like distribution of basal feet labeled with CNTRL. Scale bar represents 2 μm. **b)** 2D projection micrograph of 3DSIM volume of human airway multiciliated cell labeled with anti-CEP128 antibody. Arrows indicate hybrid cilia with ring-like distribution of basal feet labeled by anti-CEP128 antibodies. Scale bar represents 1 μm.

Figure S7. Pulse-chase experiment to track parental centrioles at the multiple basal body stage in ependymal cells

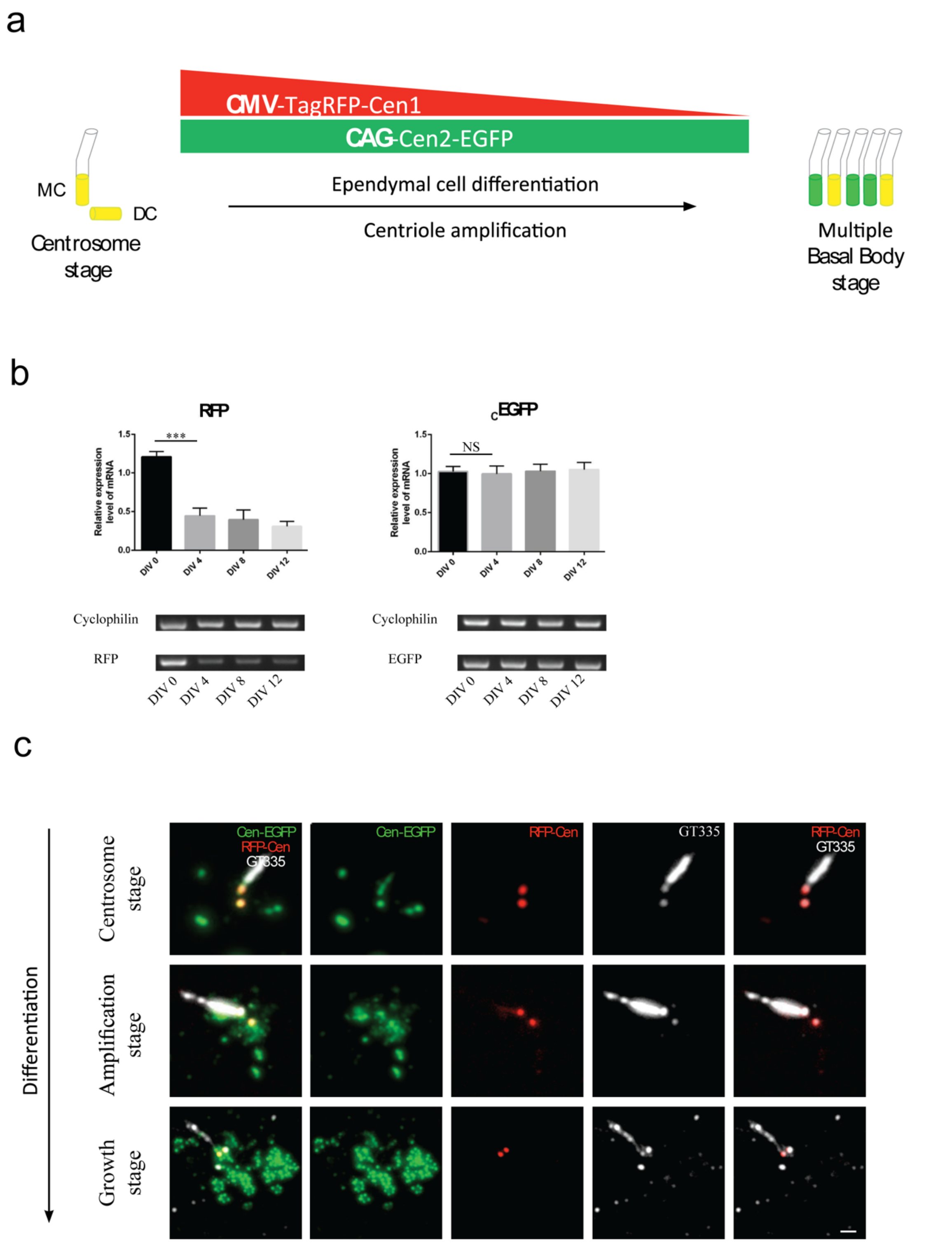

**Figure S7. Pulse-chase experiment in ependymal progenitor cells**

**a)** Cartoon depiction of the pulse-chase experiment. **b)** Semi-quantitative RT-PCR experiments show that TagRFP-Cen1 expression drops between DIV0 and DIV4. **c)** 2D projection micrograph from immunofluorescence experiment of ependymal progenitor cells labeled with TagRFP-Cen1, Cen-eGFP and anti-Glutamylated antibodies (GT335) to label cilia. Note that the different temporal expression of the markers allows a selective labelling of GT335+ centrosomal centrioles in differentiating ependymal cells.

### Figure S8: Rotational polarity and alignment vector analysis

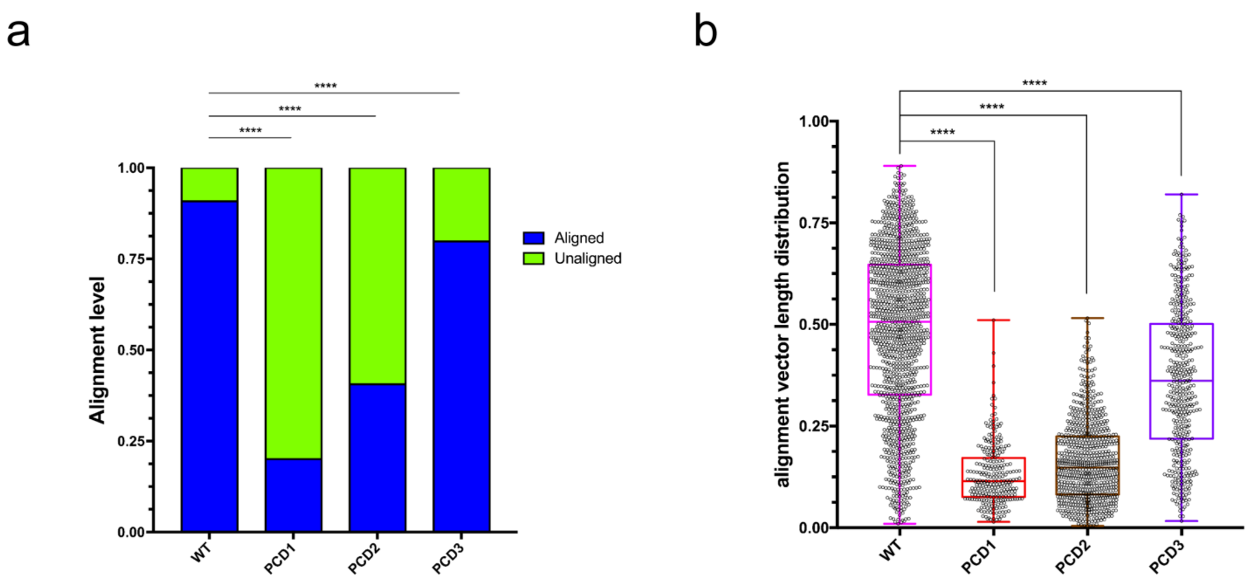


**Figure S8. Rotational polarity and alignment vector length analysis**

**a)** Stacked column graph showing the percentage of aligned (blue) and unaligned (green) cells over total cell population of WT (n=1273), PCD1 (n=274), PCD2 (n=763), and PCD3 (n=451). Statistical test was conducted using Fisher's exact test. **b)** Boxplot representing alignment vector length distribution of WT (n=1273), PCD1 (n=277), PCD2 (n=792), and PCD3 (n=453). Statistical test was conducted using one-way ANOVA and Tukey's multiple comparison test.

**Supplemental Video S1**. FIB-SEM tomograms of multiple airway multiciliated cells grown in air-liquid interface. 10 nm isotropic resolution

**Supplemental Video S2.** High magnification of Supplemental Video S1

**Supplemental Video S3.** FIB-SEM tomograms of multiple airway multiciliated cells grown in air-liquid interface. 5 nm isotropic resolution

**Supplemental Video S4.** Live imaging of TagRFP-Cen1 centrosomal centrioles during centriole amplification in primary cultured ependymal progenitors from Cen2-EFGP mice.
